# Spatial organization of neural responses to physical and agentive movement dynamics is reflected in intrinsic functional connectivity

**DOI:** 10.64898/2026.08.08.741969

**Authors:** Seda Karakose-Akbiyik, Alfonso Caramazza

## Abstract

Making sense of dynamic scenes requires interpreting the movements of inanimate objects governed by external physical forces and the actions of animate agents pursuing endogenous goals. Prior research has identified regions preferring physical or agentive movement, but their spatial organization relative to one another remains unclear. We used fMRI and within-individual analyses to examine neural responses during a motion prediction task in which two dots moved either according to physical forces (physical condition) or in coordinated, self-propelled ways suggesting intentional action (agentive condition). Resting-state data from the same participants independently characterized functional connectivity. Preferential responses to physical and agentive movement were interdigitated across frontal, parietal, and temporal cortices. Regions sharing a preference were intrinsically connected even when widely separated, while regions with opposing preferences belonged to separate networks even when adjacent. Together, these results reveal that differences between physical and agentive dynamics are not confined to local task-evoked preferences but are embedded within the brain’s broader functional organization.

## Introduction

When we look at a dynamic scene, we can effortlessly make sense of what is happening. Watching someone reach for an item on a high shelf, for instance, we not only recognize the people, objects, and the scene in which the event is taking place but also understand the interactions among them. Some of these processes apply broadly to both animate agents and inanimate objects: we register low-level visual information, track movement trajectories, and evaluate physical properties of all kinds of things in motion. Others are specific to animate agents: we interpret their intentions, anticipate their goals, and predict possible outcomes such as whether they might fall or succeed in their action. How the brain organizes these distinct aspects of dynamic scene processing, those shared across all moving entities and those specific to animate agents, is not well understood. Here, we map how responses to physically driven and agent-driven movement are arranged relative to one another within individual brains and ask whether that arrangement is reflected in intrinsic functional connectivity.

Research shows that various cortical regions across frontal, parietal, and posterior temporal lobes play a central role in interpreting dynamic scenes involving animate agents or inanimate objects. Traditionally, studies of dynamic scene processing have contrasted biological motion of animate agents with various forms of non-biological motion (e.g., object motion, random dot motion, coherent motion), often treating the latter as a baseline to isolate processes unique to animate agents (e.g., Gilaie-Dotan et al., 2013; Koldewyn et al., 2011; Saygin et al., 2004; Scholl & Gao, 2013). This line of work primarily focuses on the neural mechanisms underlying recognition and planning of human actions, typically in terms of motor representations and goals (Caspers et al., 2010; Gazzola & Keysers, 2009; Grèzes et al., 2001; Grossman & Blake, 2002; Hardwick et al., 2018; Liu et al., 2024b; Ptak et al., 2017). More recently, however, the same neural structures have been implicated in physical reasoning (Fischer et al., 2016; Schwettmann et al., 2019), and in representing the spatiotemporal structure of events more broadly (Karakose-Akbiyik et al., 2023, 2024; Wurm & Erigüç, 2025).

An emerging view holds that while frontoparietal regions involved in action, such as the premotor cortex and superior and inferior parietal lobules, constitute a domain-general system for encoding the physics and kinematics of any kind of movement (Karakose-Akbiyik et al., 2023; Kim et al., 2026), the representation of agent-specific information about dynamic scenes (e.g., biological motion, social interactions) depends on the posterior superior temporal sulcus (Deen et al., 2015, 2023; Kennedy & Adolphs, 2012; Wang et al., 2018). This region sits within a lateral occipitotemporal pathway that has recently been proposed as a “*third visual pathway*,” anatomically and functionally distinct from the classical ventral and dorsal streams and specialized for processing the dynamic visual information that supports social cognition (Pitcher & Ungerleider, 2021; Yan et al., 2025).

Not all empirical findings fit neatly within this framework, however. For instance, some studies report preferential responses to biological motion or agent movement in certain frontoparietal regions (e.g., Grèzes et al., 2001; Koldewyn et al., 2011; Ziccarelli et al., 2022) while others show preferential recruitment for inanimate object movement in overlapping regions (Beauchamp & Martin, 2007; Fischer et al., 2016). Some describe these same frontoparietal regions as the “*mirror neuron system*” (Van Overwalle & Baetens, 2009) or “*action observation network*” (Caspers et al., 2010; Hardwick et al., 2018; Urgesi et al., 2014), emphasizing the specialized role they play in making sense of the acts of conspecifics, while others focus on their general role in making sense of physical dynamics of a scene more broadly (Albertini et al., 2021; Fischer et al., 2016; Karakose-Akbiyik et al., 2023, 2024; Liu et al., 2025; Wurm & Erigüç, 2025). Furthermore, in this framework, the role of lateral occipitotemporal cortex is often underspecified, despite its consistent recruitment across paradigms that involve dynamic scene processing and converging evidence that it encodes representations of actions and event structure (Lingnau & Downing, 2015; Wurm & Caramazza, 2019).

A separate but related limitation concerns the resolution at which these questions have been studied. Most comparisons of animate and inanimate movement rely on group-averaged data, which can obscure fine-grained differences between nearby structures (see Braga & Buckner, 2017; Gordon et al., 2017). Within-subject analyses, on the other hand, typically use functional localizers to define regions of interest, then probe response properties within these predefined regions. While valuable for assessing the functional properties of specific predefined regions, this approach risks missing the broader spatial organization of neural responses, especially in neighboring areas.

Brain regions involved in dynamic scene processing have also been studied in a largely separate literature on intrinsic functional connectivity, where they are assigned to multiple large-scale association networks (Fox et al., 2005, 2006; Laufs et al., 2003; Yeo et al., 2011). For example, frontoparietal regions recruited during action planning or observation are often discussed as part of the *dorsal attention* and *frontoparietal control* networks (Ptak, 2012), which are typically studied in the context of externally directed tasks, spatial attention, and eye–hand coordination (Buschman & Kastner, 2015; Corbetta et al., 2008; Corbetta & Shulman, 2002; Dixon et al., 2018; Dosenbach et al., 2007; Heilman & Valenstein, 1972; Kastner & Ungerleider, 2000; Ptak et al., 2017; Vallar & Perani, 1987). In contrast, superior temporal and inferior frontal regions implicated in social cognition align partially with the default mode network, sometimes named the “*social brain*” (Adolphs, 2009; Amodio & Frith, 2006; Buckner & DiNicola, 2019; Lieberman, 2007; Mars et al., 2012; Meyer, 2019; Yeshurun et al., 2021). This network has broadly been associated with theory of mind (Andrews-Hanna et al., 2010b; Buckner & DiNicola, 2019; Raichle, 2015; Schurz et al., 2014).

The relationship between neural responses during dynamic scene processing and functional connectivity, however, has typically been drawn from general comparisons across paradigms or from canonical parcellations derived from group-level data (e.g., Jack et al., 2013; Simone et al., 2025). To our knowledge, it has not been directly tested within individuals. In other domains, intrinsic functional connectivity has been shown to predict the spatial layout and representational geometry of cortical response profiles, providing insights into functional organization. For example, in ventral temporal and lateral occipitotemporal cortices, connectivity structure predicts the organization of object categories, including distinctions related to animacy (Konkle & Caramazza, 2017). Extending this logic to dynamic scenes, a similar animacy-related organization in higher-level regions may be expressed not through category selectivity per se, but through sensitivity to differences in how motion is generated, specifically, whether movement is driven by external physical forces or by intentional agents.

In this study, we explore animacy as a candidate organizing principle in how the brain processes information about dynamic scenes. Specifically, we focus on one aspect of animacy that reflects variation in kinematic patterns: movements driven solely by external physical forces (physical movement) versus those generated by intentional agents (agentive movement). Motivated by the limitations of group-level and localizer-based approaches, we adopt a within-subject analysis strategy that can characterize the spatial organization of neural responses across neighboring regions rather than within predefined areas.

Our objectives were twofold: (1) to characterize the topography of distributed neural responses to physical and agentive movement within individuals, and (2) to determine the degree to which the spatial organization of preferential responses to physical or agentive movement is reflected in intrinsic network architecture. We find that preferential responses to physical and agentive movement dynamics are not confined to anatomically distant regions but frequently interleave across closely neighboring regions of frontal, parietal, and temporal cortex. Mapping these response preferences onto individualized functional networks further reveals that this fine-grained spatial organization aligns with the brain’s intrinsic network architecture. Together, our results show that the distinctions between physical and agentive movement dynamics are reflected not only in local response preferences but also in the distributed organization of large-scale brain networks.

## Results

### Procedure

To situate neural responses to physical and agentive movement within the context of intrinsic functional connectivity, we conducted two scanning sessions per participant (n = 12), each including task runs and resting-state runs. The experimental task was a motion prediction task (DOTS task) in which participants viewed movies of two moving dots (Fischer et al., 2016). The trajectories of the dots represented two motion categories defined here as physical (i.e., determined by Newtonian mechanics for collisions) and agentive (i.e., self-propelled movement depicting agent-like interactions, see Figure 1A).

**Figure 1.**
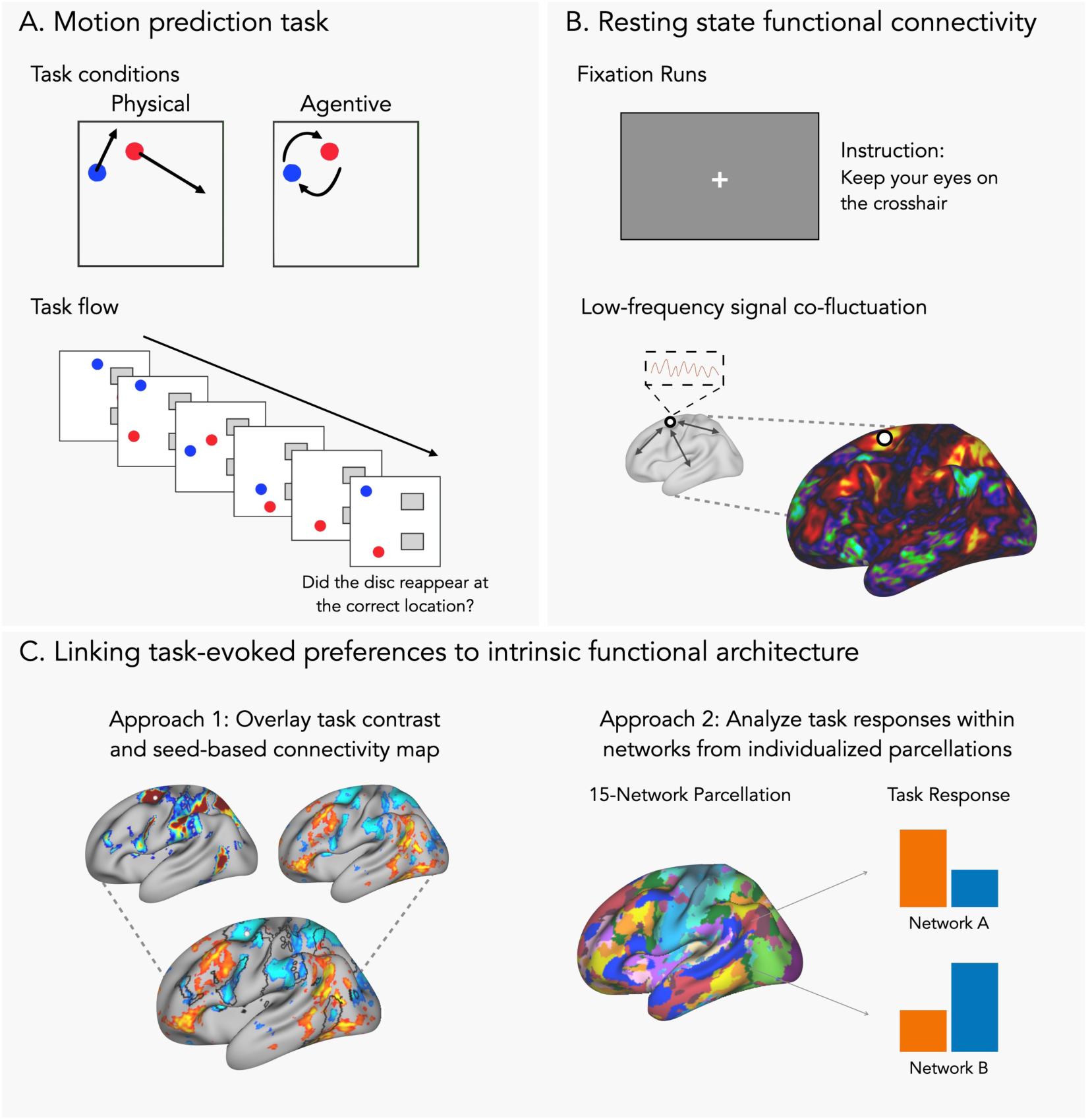
Overview of the fMRI paradigms and analysis pipeline. **(A) Motion prediction task.** Participants viewed movies depicting interactions between two dots, defined here as physical or agentive. During the movie, the dots moved around until one dot disappeared and reappeared in a new location. Participants judged whether the dot reappeared in the correct location by pressing a button. The icons representing physical and agentive movement in the figure were adapted from Liu et al. (2024a). **(B) Resting-state functional connectivity.** Participants completed resting-state fixation runs, maintaining gaze on a central crosshair without performing any task. Low-frequency signal co-fluctuations were measured to identify regions with correlated spontaneous activity. **(C) Linking task-evoked preferences to intrinsic functional architecture.** Two approaches were used to examine the relationship between task-evoked preferences and network organization. In Approach 1, we placed seeds in regions showing preferential responses to physical or agentive movement to qualitatively examine the spatial correspondence between task activations and connectivity-derived networks. In Approach 2, functional response dissociations were quantified within specific networks defined by individualized 15-network parcellations (MS-HBM, see Du et al., 2024). Average responses to physical and agentive movement were examined within each network.

For both types of stimuli, the dots moved around for eight seconds before one became invisible for two seconds and then reappeared. Participants were asked to imagine the trajectory of the invisible dot and judge whether it reappeared at the correct location. In the physical condition, this judgment can be made solely based on physical properties of movement, for example, predicting how an object’s trajectory will change following a collision. The agentive condition, on the other hand, requires inferring the agents’ intentions, for example, recognizing that one agent is chasing another and predicting its movement accordingly.

Research using this and similar paradigms has largely adopted a contrastive approach, focusing on identifying regions of interest that show differential activity between categories, often based on relatively limited data per participant. While this approach is useful in identifying regions of interest, it is less suited for characterizing the complex, distributed nature of neural responses. Given our goal of examining the topography of neural responses within individuals, we collected approximately two to three times the typical amount of data per participant to improve signal-to-noise and enable precise characterization of within-individual response patterns (see Methods for more detail).

For the analyses of intrinsic functional connectivity, and to situate neural responses in the DOTS task within the functional network architecture of the brain, participants also completed resting-state fixation runs in which they kept their eyes on a fixation cross without performing any specific task (see Figure 1B). Low-frequency signal co-fluctuations were used to identify regions with correlated spontaneous activity. We collected at least thirty-five minutes of resting-state data per participant, allowing individualized analyses of intrinsic functional connectivity through seed-based and parcellation-based approaches (see Du et al., 2024; Kong et al., 2019). As our analyses focused on within-individual organization, we used a preprocessing pipeline optimized for cross-session alignment and minimal spatial blurring (Braga et al., 2019; Du et al., 2024; see Methods for more details on the preprocessing of the fMRI data).

### Distributed cortical regions respond to physical and agentive movement

The DOTS task formed the basis for examining neural responses to physical and agentive movement. To identify the brain regions recruited in each condition, we first visualized brain regions showing increased responses against baseline for both conditions. Figure 2 illustrates the recruitment profiles for the physical and agentive conditions against baseline, as well as their overlap, for sample individuals (*p* < .0001, *z* > 3.72, one-tailed, uncorrected; see Supplementary Figure 1 for all participants). Blue indicates regions showing higher-than-baseline activity selectively for physical movement, whereas red indicates regions showing higher-than-baseline activity for agentive stimuli. Purple denotes regions that showed above-baseline activity for both conditions.

**Figure 2.**
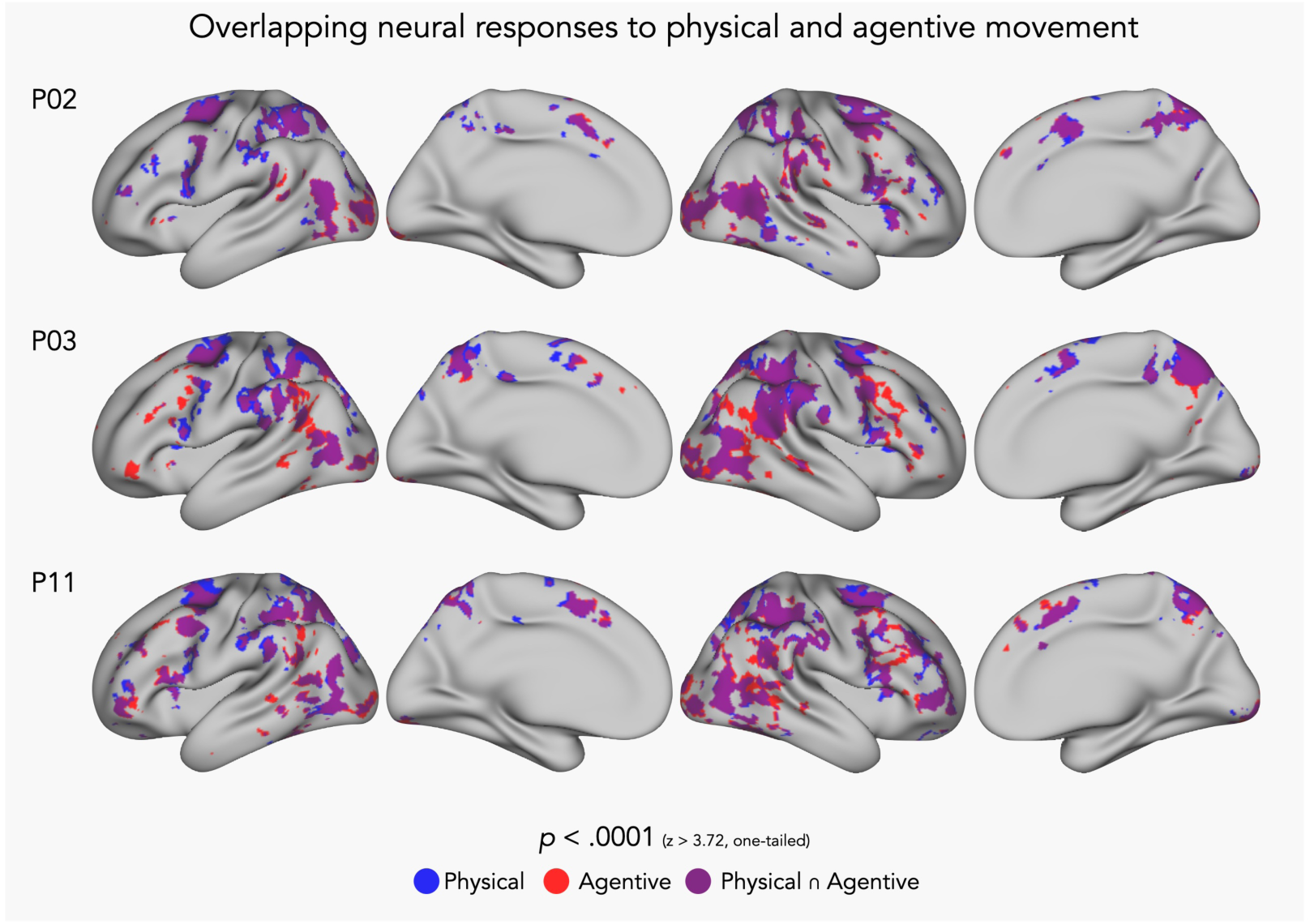
Neural responses to physical and agentive movement in sample participants. Cortical regions with increased responses compared to baseline during the DOTS task for physical and agentive movement. Blue regions represent regions with higher than baseline activity exclusively for physical movement, red regions for agentive movement, and purple regions show the overlap where both conditions elicited higher than baseline activity. Threshold: *p* < .0001, z > 3.72, one-tailed, uncorrected.

As shown in Figure 2, neural responses to physical and agentive movement against baseline largely overlapped. To quantify spatial overlap between suprathreshold vertices (thresholded at *p* < .0001 uncorrected) in the two conditions, we computed the Dice coefficient. The Dice coefficient was defined as

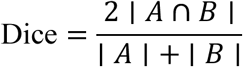

where A and B represent the sets of suprathreshold vertices in the two conditions. Dice coefficients averaged 0.84 ± 0.04 (mean ± SD), indicating strong overlap of above-baseline activity between physical and agentive movement. Both conditions activated parts of the dorsal and ventral premotor cortex, superior and inferior parietal lobule, and lateral occipitotemporal cortex, bilaterally. Notably, across all subjects, regions showing above-baseline responses exclusively for the physical condition (blue) were limited. However, several regions showed higher-than-baseline activity only in the agentive condition (red), particularly around posterior superior temporal sulcus, dorsolateral prefrontal cortex, and ventral premotor cortex, especially in the right hemisphere (see Supplementary Figure 1 for data from all subjects).

Overlapping recruitment for these two conditions relative to baseline is expected, as both involve analyzing and predicting the kinematic patterns of moving dots. Importantly, the baseline consisted of an empty screen, such that above-baseline activity likely reflects general visual and cognitive engagement common to both conditions rather than processes specific to physical or agentive dynamics. Accordingly, we do not interpret the individual condition-versus-baseline maps in detail but instead present them as context for identifying where the conditions diverge, which we examine next.

### Preferential responses to physical and agentive movement are distributed and spatially interleaved

Next, we examined the large-scale spatial organization of preferential neural responses to physical or agentive movement by comparing the two conditions. Based on previous literature that used the same paradigm, we expected preferential responses in regions spanning the premotor cortex, superior parietal lobule, and inferior parietal lobule for the physical condition, and in the right pSTS for the agentive condition (Fischer et al., 2016; Liu et al., 2024a).

To explore distributed preferences for physical versus agentive stimuli, we used a liberal threshold for the physical-agentive contrast map at *p* < .01 (|z| > 2.58, two-tailed, see Figure 3). We provide maps from all participants at *p* < .05 (|z| > 1.96) to convey the full range of individual variability (Supplementary Figure 2). These maps are used for visualization of distributed trends only; none of the statistical inferences reported below depend on them as all quantitative tests were conducted on network-level averages rather than on thresholded vertices.

**Figure 3.**
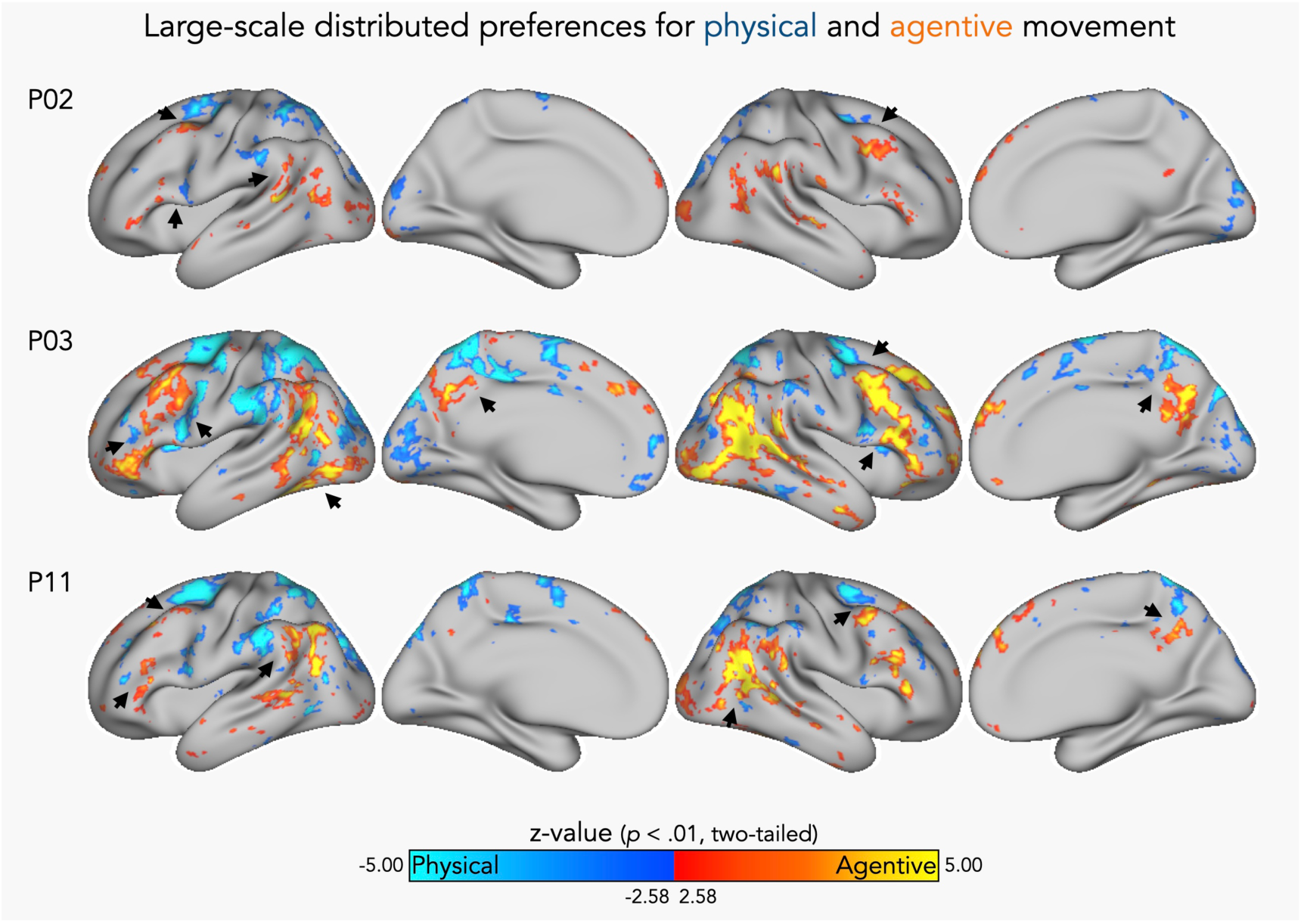
Univariate contrast of physical and agentive movement in sample participants. Neural response differences between physical and agentive conditions. Blue regions show greater responses to physical stimuli, while orange/yellow regions show greater responses to agentive stimuli. Threshold: *p* < .01, |z| > 2.58, two-tailed, uncorrected. Arrows indicate anatomically adjacent clusters with distinct preference profiles, highlighted for illustrative purposes.

Across participants, preferential responses to physical movement were observed around dorsal and ventral premotor cortex, superior and anterior inferior parietal lobule, and, in some participants, anterior lateral occipitotemporal cortex (see Supplementary Figure 2 for all participants). In contrast, preferential responses to the agentive condition mostly occurred around the posterior superior temporal sulcus (pSTS), and temporoparietal junction (TPJ), but also extended into additional regions in the frontal cortex (ventral premotor cortex and dorsolateral and ventrolateral prefrontal cortex). Note that the spatial topography of physical-preferring responses was more reproducible across participants with largely consistent loci, whereas agentive-preferring responses exhibited greater inter-individual variability (see Supplementary Figure 2).

Visual inspection of individual activation maps showed that neighboring regions across frontal, parietal, and temporal cortices often exhibited distinct preference profiles (see Figure 3). Overall, preferences for physical and agentive movement were not confined to isolated loci but were distributed across multiple cortical regions, raising the question of whether regions with a similar response profile belong to a common functional network.

### Distributed cortical preferences for physical and agentive movement are reflected in intrinsic functional connectivity

After exploring the distributed nature of neural responses to physical and agentive movement, we assessed whether spatially distributed regions that preferentially respond to one type of movement constitute a unified network of interconnected regions at rest. To situate neural responses in the motion prediction task within the context of intrinsic functional connectivity, we used both seed-based and parcellation-based approaches (see Figure 1C).

First, we tested the relationship between task responses and intrinsic functional connectivity using a seed-based approach. We began by identifying candidate anatomical loci for each contrast (physical > agentive and agentive > physical) based on cortical regions that showed consistent preferential responses across participants. For each participant, we then selected specific seed vertices within these loci that resulted in a network with strong correlations with the seed region (*z*(r) ≈ 0.60). Seed selection prioritized locations whose network organization aligned with the broader task response rather than simply the voxel with the highest activation. To test the spatial correspondence between intrinsic connectivity patterns and task activations, we overlaid the resulting connectivity maps with task responses (see Figure 4 for the physical seed, and Figure 5 for the agentive seed).

**Figure 4.**
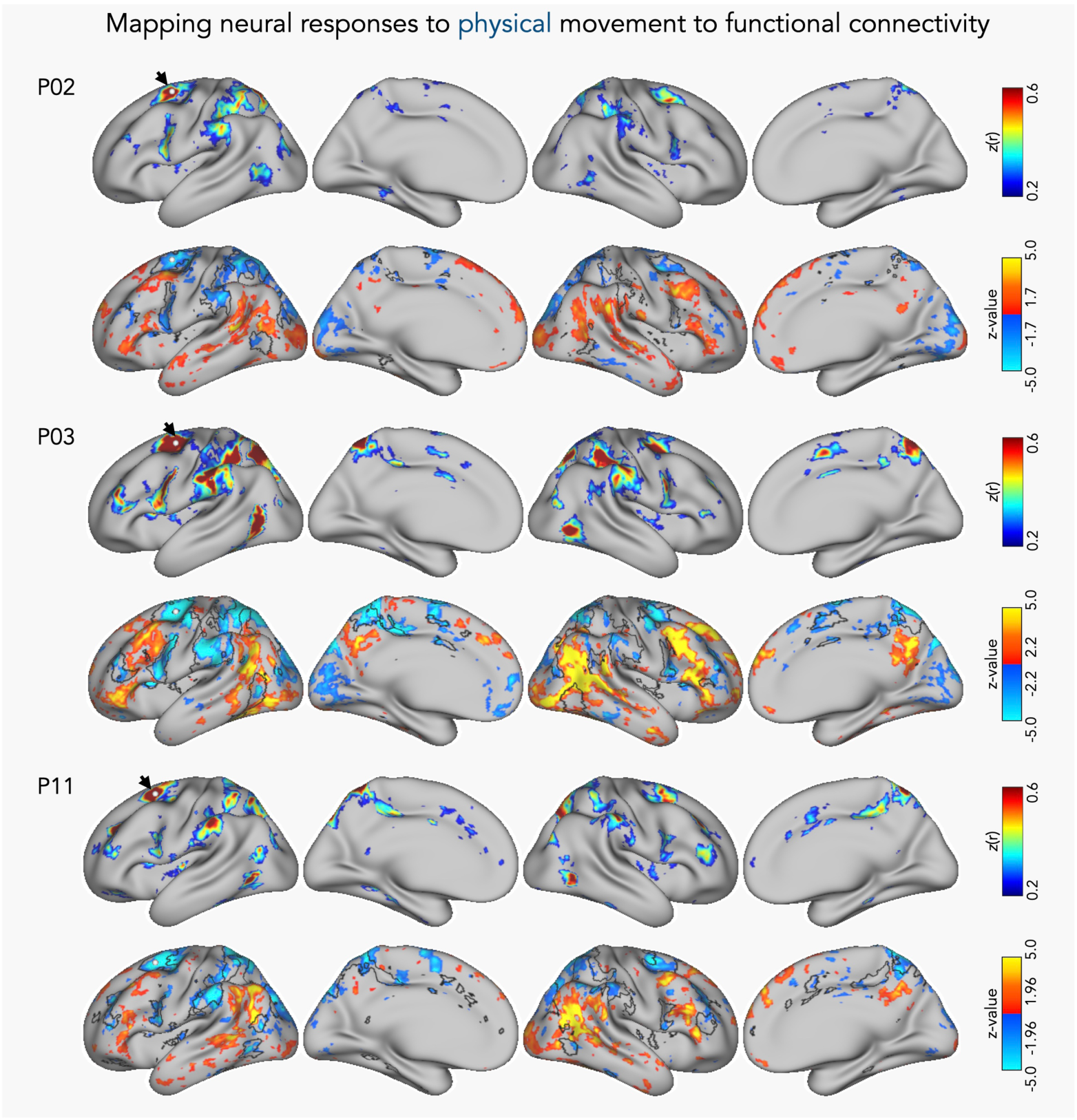
Mapping preferential responses to physical movement to intrinsic functional connectivity. For each participant, the top panel shows the seed-based functional connectivity map generated from the physical-condition seed, with the white dot marking the seed location highlighted by a black arrow for visibility. The functional connectivity maps are thresholded with *z*(r) values between 0.20 and 0.60. The bottom panel shows the DOTS task contrast, with regions preferring physical (blue) and agentive (orange/yellow) movement, overlaid with the outline of the corresponding functional connectivity map. For visualization purposes, contrast maps were displayed using a liberal and varying threshold (*|z|* = 1.96-2.20) to better illustrate individual trends.

**Figure 5.**
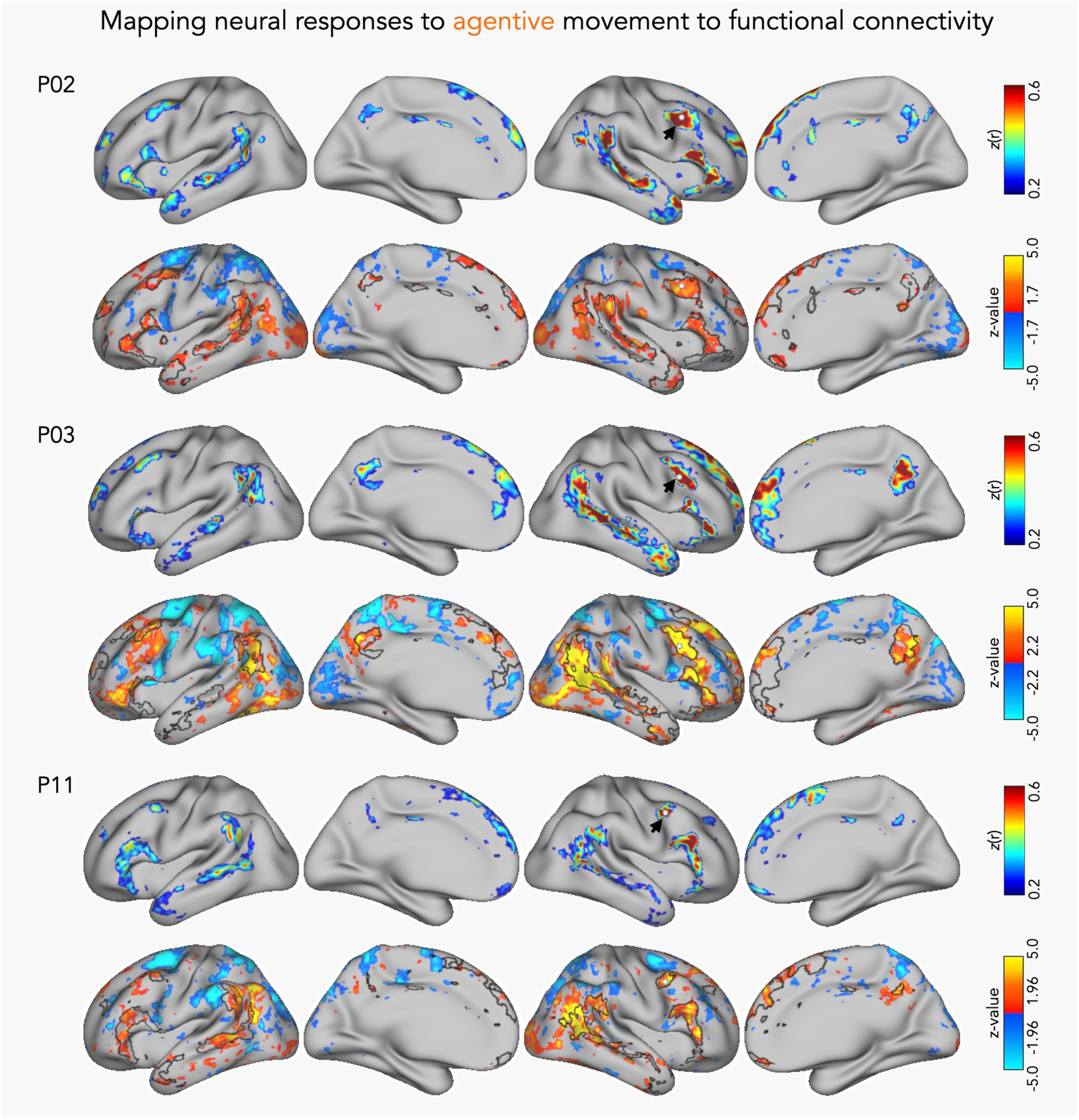
Mapping preferential responses to agentive movement to intrinsic functional connectivity. For each participant, the top panel shows the seed-based functional connectivity map generated from the agentive-condition seed, with the white dot marking the seed location highlighted by a black arrow for visibility. The functional connectivity maps are thresholded with z(r) values between 0.20 and 0.60. The bottom panel shows the DOTS task contrast, with regions preferring physical (blue) and agentive (orange/yellow) movement, overlaid with the outline of the corresponding functional connectivity map. For visualization purposes, contrast maps were displayed using a liberal and varying threshold (*|z|* = 1.96-2.20) to better illustrate individual trends.

In ten out of twelve subjects, the peak for the physical > agentive contrast was found around the left dorsal premotor cortex. Accordingly, we identified seeds in this region for testing the spatial correspondence between functional connectivity and preferential responses to the physical condition for each individual subject. The dorsal premotor cortex includes components that are functionally coupled with the canonical dorsal attention system; therefore, we expected the connectivity of this seed to reveal a network overlapping with regions of the so-called dorsal attention system (Kong et al., 2019; Yeo et al., 2011).

Figure 4 shows the connectivity profiles of the seed selected for the physical condition and its overlay with the corresponding task contrast for sample participants (see Supplementary Figures 3-4 for all participants). The white dot marks the location of the seed used for the connectivity analysis. The connectivity maps are thresholded at *z*(r) = 0.20 to visualize the resulting networks (Braga & Buckner, 2017). For visualization, task contrast maps are displayed using a liberal and varying threshold to better illustrate individual trends (for a similar approach, see DiNicola et al., 2020; Du et al., 2024).

The connectivity profile of the dorsal premotor seed revealed a bilateral network encompassing the premotor cortex, superior and inferior parietal lobules, and anterior lateral occipitotemporal cortex. To test the spatial correspondence of this network with the neural response preferences observed in the DOTS task, we overlaid the outline of the connectivity map with task-based contrast maps. As shown in Figure 4, overlaying this network with the task contrast maps revealed that regions preferring physical movement largely fell within the network defined by the dorsal premotor seed, with clear boundaries separating them from neighboring agent-preferring regions, in both hemispheres. These abutting patterns were especially evident near premotor cortex where side-by-side regions displayed distinct preference profiles (for all participants, see Supplementary Figure 4).

Notably, a region in anterior lateral occipitotemporal cortex (LOTC) consistently appeared as part of the network defined by the dorsal premotor seed, despite not showing a reliable preferential response to physical over agentive movement in the univariate contrast. Besides its established role in action recognition (Lingnau & Downing, 2015), LOTC is also commonly recruited in functional neuroimaging studies on physical reasoning. For example, localizers contrasting judgments about whether a block tower will fall with judgments about its color identify preferential responses in LOTC alongside frontoparietal regions typically described as the “physics network” (see Fischer et al., 2016; Schwettmann et al., 2019).

In recent literature on physical reasoning, however, LOTC is often analyzed separately from frontoparietal regions. Some studies suggest that frontoparietal regions but not LOTC encode physical properties in a more abstract and generalizable manner (Pramod et al., 2022; Schwettmann et al., 2019). These studies often define LOTC using relatively broad anatomical regions, potentially obscuring finer-grained distinctions within this heterogeneous territory. The present results argue against treating LOTC as a single functional unit.

A cluster in anterior LOTC was functionally coupled with the frontoparietal regions implicated in processing physical scene dynamics, whereas posterior LOTC fell outside this network and instead showed clusters with preferential responses to agentive movement in some participants. Although a few participants had small physical-preferring clusters in anterior LOTC, the region as a whole did not reliably prefer either condition despite being embedded in the network identified by the physical seed. This dissociation between intrinsic connectivity and univariate response preference suggests that the anterior LOTC cluster may play a role within this network that is not captured by the physical–agentive contrast.

We next applied the same approach to examine the relationship between intrinsic functional connectivity and preferential responses to agentive movement (see Figure 5). Although preferential responses to the agentive condition were most strongly and consistently observed around the right pSTS, we placed seeds at or near the ventral premotor cortex, at the intersection of the posterior middle frontal gyrus and precentral sulcus. This location was selected because ten out of twelve participants showed clusters with a preference for the agentive condition in this region, providing a consistent anatomical basis for comparison with the physical-condition seed in the dorsal premotor cortex. Moreover, at or near premotor cortex, adjacent vertices frequently showed alternating preferences for the physical or agentive condition (see Supplementary Figure 2). Demonstrating that these spatially nearby vertices are embedded within distinct intrinsic networks would provide particularly strong evidence that the fine-grained organization of task-evoked responses reflects the brain’s intrinsic functional architecture.

Figure 5 shows the connectivity profile of the seeds selected for the agentive condition and their overlay with the corresponding task contrast for sample participants (see Supplementary Figures 5 and 6 for all participants). The seed selected for the agentive condition revealed a different bilateral network encompassing temporoparietal junction, superior temporal sulci, portions of lateral prefrontal cortex, and posterior medial regions. Thus, even though this network included the TPJ and pSTS that are commonly associated with processing of theory of mind, biological motion, and social interactions, it also extended into various other frontoparietal loci, often forming interdigitated patterns with regions preferring physical movement.

**Figure 6.**
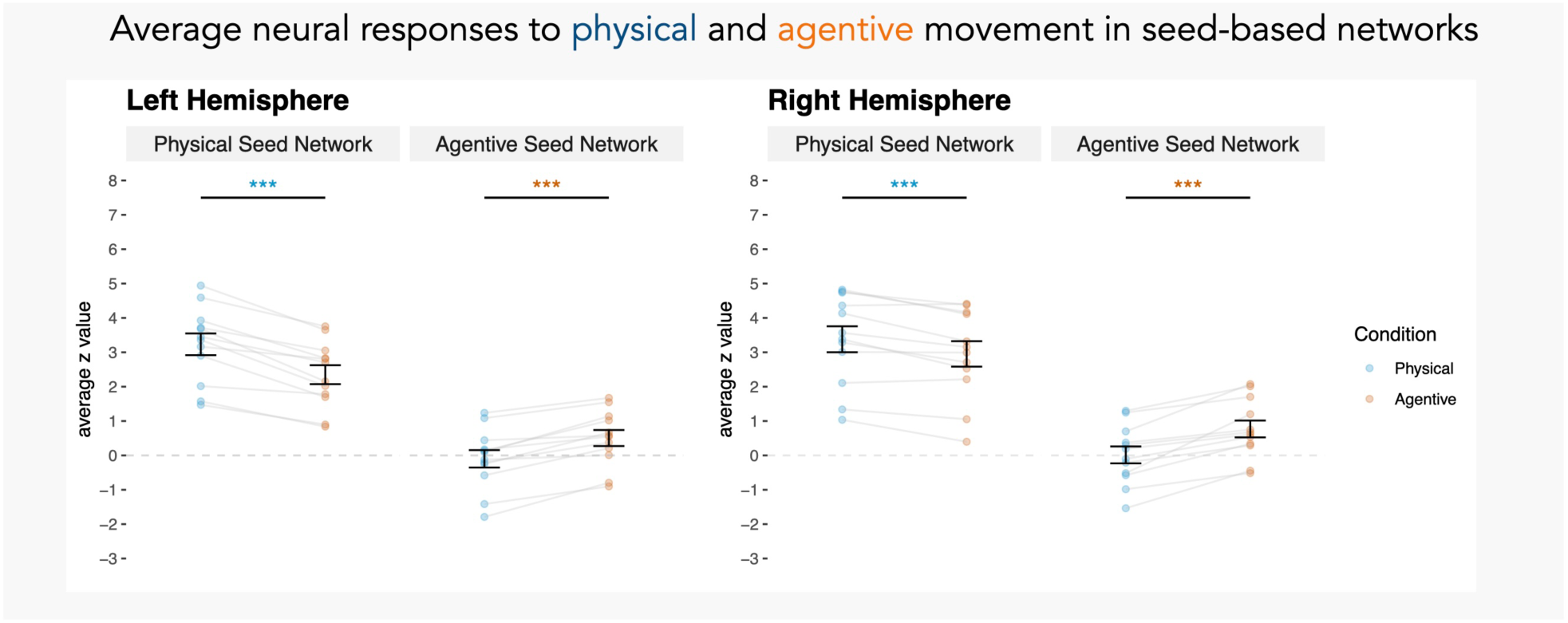
Average neural responses to the agentive and physical conditions within networks estimated by seed-based functional connectivity. Significant differences are marked and colored by the condition that showed the higher average response (blue for physical, orange for agentive; FDR-corrected, \**p* < .05, \*\**p* < .01, \*\*\**p* < .001). Error bars indicate SEM.

Note that preferential responses to agentive movement were more variable across participants, showing individual differences in both the task maps and the corresponding networks based on functional connectivity. For some subjects, we did not find clusters with preference for the agentive condition near premotor cortex, also making it difficult to select an appropriate seed to align with task responses (e.g., P6 and P9 in Supplementary Figure 6). Furthermore, unlike the physical condition where the network outline encompassed nearly all frontoparietal and temporal regions showing preference for physical movement, the network derived from the agentive seed did not always align with task responses in every participant.

For instance, in several participants, the agentive condition elicited preferential responses in parts of posterior lateral occipitotemporal cortex that frequently fell outside the boundaries of the network derived from the prefrontal seed (e.g., see P5 and P9 in Supplementary Figure 6). This is plausible given that the two stimulus conditions differed along multiple dimensions (e.g., low-level kinematics, interaction structure, and the requirement to infer goals or social contingencies). We think that the network identified by the agentive seed likely indexes recruitment of higher-level interpretive processes related to the agentive condition (intent/social inference), which may be engaged to varying degrees across participants depending on what they paid attention to and task strategy. In contrast, agentive-preferring responses in the posterior occipitotemporal foci likely reflect sensitivity to visual/biological-motion features and other lower-level distinctions between physical and agentive interactions depicted by the stimuli. Future work that explicitly manipulates visual features of the stimuli, attentional demands, and task goals will be needed to test how these factors shape the recruitment of these partially dissociable response profiles.

Although the spatial overlap between connectivity-derived networks and task-based preferences was less consistent for the agentive than the physical condition, the resulting networks rarely, if ever, included regions preferring physical movement. Notably, similar to the physical condition, network boundaries often delineated anatomically close loci showing distinct preferences for physical or agentive movement. The spatial correspondence between connectivity maps and task preferences revealed that some response patterns that might initially appear noisy were, in fact, highly systematic, closely following network boundaries (see Figures 4 and 5). In some regions, particularly around the frontal and parietal cortices, vertices showing opposing response preferences (physical vs. agentive) were often situated side by side, yet their arrangement mirrored the fine-grained organization of the corresponding functional connectivity patterns. This spatial correspondence suggests that the interdigitated response patterns are systematically related to intrinsic functional network organization rather than reflecting arbitrary spatial variation.

To quantify the responses of the networks identified through seed-based correlations, we treated each network as a region of interest. First, we assessed the consistency of vertex-wise responses by quantifying the proportion of vertices exhibiting the expected condition preference. Within networks defined by the physical seed, the majority of vertices exhibited higher responses to the physical than to the agentive condition (LH: 79.90% ± 11.00, RH: 69.60% ± 14.50). Conversely, within networks defined by the agentive seed, the majority of vertices exhibited higher responses to the agentive than to the physical condition (LH: 77.10% ± 13.10, RH: 80.20% ± 9.63). Together, these patterns indicate that the response preference was distributed across most of each network rather than restricted to a subset of vertices.

We also quantified the mean response magnitude within the boundaries of each respective network, allowing us to evaluate how strongly these networks responded to the physical and agentive conditions, respectively. To test for differences between neural responses to physical and agentive movement across these seed-based networks, we computed a two-way repeated-measures ANOVA for each hemisphere testing the interaction between network and condition. Following significant interactions, we conducted pairwise t-tests with p values corrected for multiple comparisons using the false discovery rate (FDR).

A repeated-measures ANOVA revealed a significant network × condition interaction in both hemispheres (LH: *F*(1, 11) = 63.44, *p* < .001, η²₍G₎ = .15; RH: *F*(1, 11) = 44.38, *p* < .001, η²₍G₎ = .07). In the left hemisphere, responses within the physical network were significantly stronger for the physical than the agentive condition (*t*(11) = 7.51, *p* < .001, *d* = 2.17, 95% CI [0.63, 1.14]), whereas responses within the agentive network were significantly stronger for the agentive than the physical condition (*t*(11) = −6.63, *p* < .001, *d* = −1.91, 95% CI [−0.81, −0.40]). A similar pattern was observed in the right hemisphere. Responses within the physical network were significantly stronger for the physical than the agentive condition (*t*(11) = 4.44, *p* = .001, *d* = 1.28, 95% CI [0.22, 0.64]), whereas responses within the agentive network were significantly stronger for the agentive than the physical condition (*t*(11) = −5.99, *p* < .001, *d* = −1.73, 95% CI [−1.03, −0.48]).

Note that although the extent of each network was determined entirely by resting-state connectivity, the seed locations were chosen with reference to the task maps. Some bias toward the defining condition is therefore expected, and these comparisons should be read as quantifying the spatial correspondence apparent in Figures 4 and 5 rather than as an independent test of it. The parcellation-based analyses reported below provide such a test.

### Parcellation-based analyses of task responses for an independent evaluation

Thus far, we have examined the correspondence between seed-based networks and task-evoked responses to physical or agentive movement through qualitative boundary comparisons and quantitative analyses of average responses within seed-based networks. However, seed-based approaches have limitations. Seed selection can introduce biases, as a given seed may not optimally capture the full extent or underlying correlation pattern of a network. This issue was particularly relevant for responses to agentive movement, which showed greater inter-individual variability in their spatial organization, and therefore in the loci used to define seeds for connectivity analyses. To conduct an independent investigation of functional network involvement in response to physical and agentive movement, and to link our findings to the broader literature on functional connectivity, we also adopted a data-driven model-based parcellation approach.

To directly test functional network involvement in the analysis of physical and agentive movement, we estimated the cortical parcellations within individual subjects using a multi-session hierarchical Bayesian model (MS-HBM, Kong et al., 2019), with a recent 15-network prior developed by Du and colleagues (Du et al., 2024). This method provides precise network estimates for each individual and distinguishes multiple networks, including parallel networks within the dorsal attention and default mode systems that were of interest for the current study (see Figure 7 for the MS-HBM group prior and sample individual parcellations, see Supplementary Figure 7 for parcellations of all participants).

**Figure 7.**
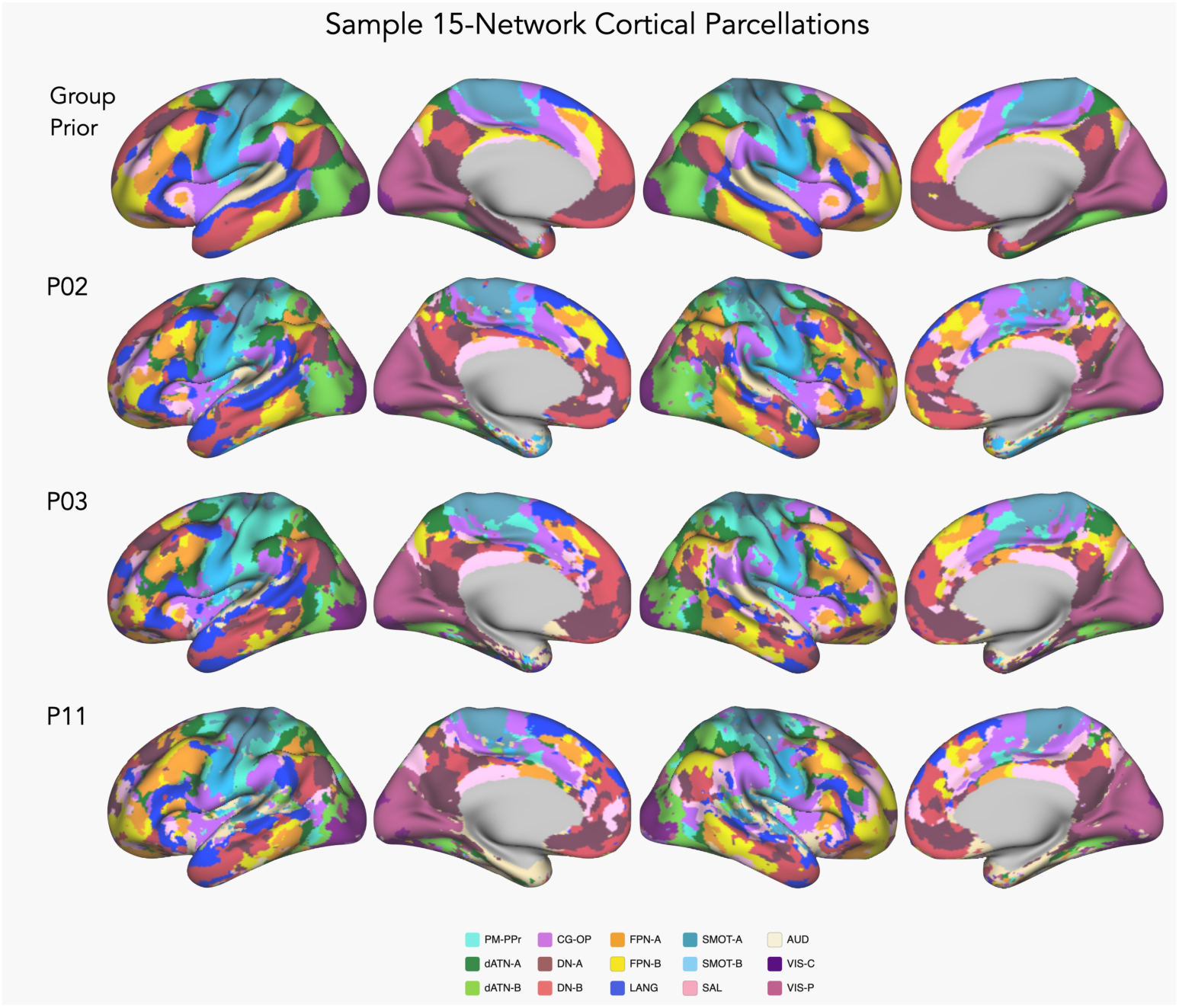
15-network parcellation group prior and sample participants. 15-network estimation for identifying the cortical networks for sample participants. Cortical networks were estimated for each participant using a multi-session hierarchical Bayesian model, following the implementation described in Du et al. (2024). The model was initialized with a group prior generated from the Human Connectome Project S900 (top row).

We focus our analyses on a subset of the 15 networks that are characterized by the model: Premotor-Posterior Parietal Rostral (PM-PPr), Dorsal Attention-A (dATN-A), Dorsal Attention-B (dATN-B), Cingulo-Opercular (CG-OP), Frontoparietal Network-A (FPN-A), Frontoparietal Network-B (FPN-B), Default Network-A (DN-A), Default Network-B (DN-B), and Language (LANG). We chose these networks because they lie outside the primary sensorimotor processing streams, have similar distributed zones across frontal, parietal and temporal cortices, and have been linked to processes relevant for interpreting dynamic scenes.

The overlap of the dorsal attention system with brain regions involved in action recognition and dynamic scene processing has been noted (Schurz et al., 2020), although not tested directly within individuals. Furthermore, while the literature often refers to a single dorsal attention system, it can be divided into three distinct networks: PM-PPr, dATN-A, and dATN-B. Each of these networks shows unique functional connectivity patterns and may exhibit distinct functional properties (Braga & Buckner, 2017). Additionally, previous work on the network basis of action and physical inference, as well as the literature on functional connectivity, often conflates frontoparietal control networks (FPN-A, FPN-B) and the dorsal attention system (PM-PPr, dATN-A, dATN-B), highlighting the need to examine these networks separately within individuals. Finally, the cingulate cortex that is encompassed by the Cingulo-Opercular network (CG-OP) is involved in action planning (Badke D’Andrea et al., 2025; Rushworth et al., 2004), shows increased functional coupling with regions involved in physical inference (Navarro-Cebrián & Fischer, 2022), and lies in close proximity to the dorsal attention system. Thus, we also included this network in our analysis.

The default mode network is of particular interest in the present context, as it has been extensively studied in relation to internally directed cognition, theory of mind, and social reasoning (Andrews-Hanna et al., 2010a). More recent work, however, has shown that the default mode network is not functionally homogeneous but can be subdivided into parallel networks that differentially support spatial and social reasoning (Deen & Freiwald, 2025; DiNicola et al., 2020). Given this distinction, the physical and agentive conditions in the DOTS task could engage these subnetworks differently: the physical condition emphasizes spatiotemporal prediction and tracking contacts with landmarks and other objects, while the agentive condition emphasizes interpreting interactions between two agents, including their social goals. Accordingly, we tested whether default mode network subcomponents linked to spatial versus social reasoning differentially track the physical and agentive conditions within individuals. Finally, the LANG network refers to the functional connectivity counterpart of the task-defined language network typically associated with lexico-semantic and syntactic processing (Braga et al., 2020; Fedorenko et al., 2024), whose regions lie in close spatial proximity to the neighboring default mode networks. Thus, we also tested whether its response profile converges with, or dissociates from, default mode networks in tracking physical or agentive movement dynamics.

To directly compare the functional network recruitment in response to physical and agentive movement dynamics, for each participant, we calculated the average z-value across vertices within each candidate network, separated by hemisphere (see Figure 8). In both hemispheres, PM-PPr, dATN-A, and dATN-B exhibited increased recruitment compared to the baseline. Conversely, DN-A and DN-B did not show increased responses against the baseline; instead, in line with the literature, they demonstrated a trend towards lower-than-baseline responses (Buckner et al., 2008; Raichle et al., 2001; Smith et al., 2018).

**Figure 8.**
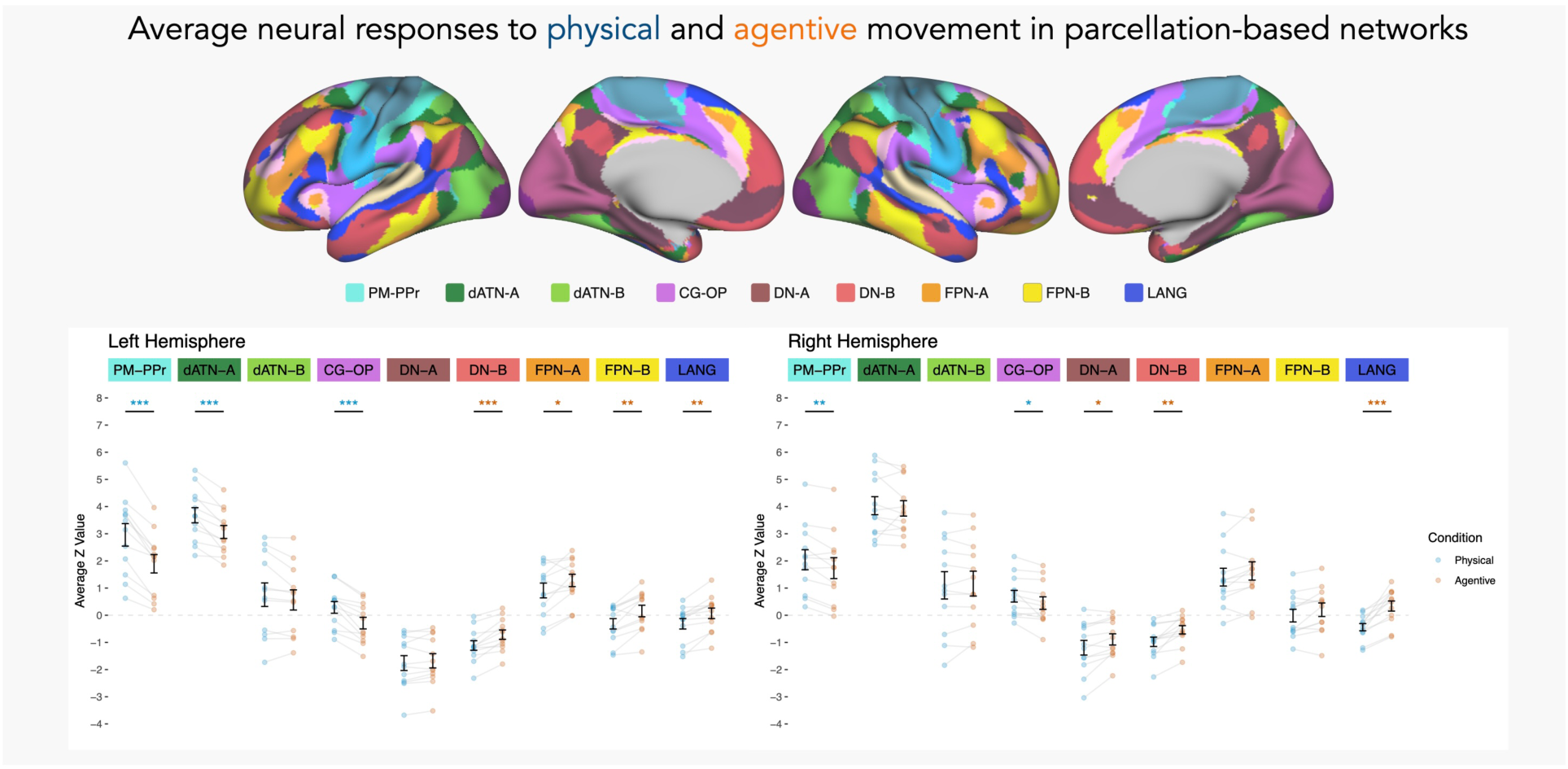
Average neural responses to physical and agentive movement by networks derived from individual parcellations. Average z-values of neural responses for physical (blue) and agentive (orange) conditions across different networks in the left and right hemispheres. Dots represent individual responses, and faint gray lines connect responses from the same individual. Significant differences are marked and colored by the condition that showed the higher average response (blue for physical, orange for agentive; FDR-corrected, \**p* < .05, \*\**p* < .01, \*\*\**p* < .001). Error bars indicate SEM.

To test for differences between neural responses to physical and agentive movement across different functional networks, we computed a repeated-measures ANOVA for each hemisphere testing the interaction between network and condition. Following significant interactions, we conducted pairwise t-tests with p values corrected for multiple comparisons using the FDR. The interaction between network and condition was significant for both left (*F*(3.03, 33.38) = 28.51, *p* < .001, η²₍G₎ = .08) and right hemispheres (*F*(3.40, 37.36) = 12.59, *p* < .001, η²₍G₎ = .03), with Greenhouse-Geisser corrected degrees of freedom and p values, indicating that the degree of difference between the physical and agentive conditions varied across networks.

For the left hemisphere, physical movement led to greater recruitment than agentive movement in PM-PPr (*t*(11) = 8.10, *p* < .001, *d* = 2.34, 95% CI [0.78, 1.35]), dATN-A (*t*(11) = 5.58, *p* < .001, *d* = 1.61, 95% CI [0.37, 0.86]), and CG-OP (*t*(11) = 6.08, *p* < .001, *d* = 1.75, 95% CI [0.37, 0.79]). An adjacent network, dATN-B, did not show a significant difference between the two conditions (*t*(11) = 1.88, *p* = .097, *d* = 0.54, 95% CI [−0.03, 0.42]). Conversely, agentive movement led to greater recruitment in DN-B (*t*(11) = −5.41, *p* < .001, *d* = −1.56, 95% CI [−0.56, −0.24]), FPN-A (*t*(11) = −2.67, *p* = .028, *d* = −0.77, 95% CI [−0.68, −0.07]), FPN-B (*t*(11) = −4.57, *p* = .001, *d* = −1.32, 95% CI [−0.70, −0.24]), and LANG (*t*(11) = −4.72, *p* = .001, *d* = −1.36, 95% CI [−0.56, −0.21]). DN-A showed no significant difference between the two conditions (*t*(11) = −1.51, *p* = .159, *d* = −0.44, 95% CI [−0.21, 0.04]).

For the right hemisphere, physical movement led to greater recruitment than agentive movement in PM-PPr (*t*(11) = 3.91, *p* = .007, *d* = 1.13, 95% CI [0.14, 0.48]) and CG-OP (*t*(11) = 2.98, *p* = .023, *d* = 0.86, 95% CI [0.07, 0.44]). Neither dATN-A (*t*(11) = 0.68, *p* = .508, *d* = 0.20, 95% CI [−0.22, 0.42]) nor dATN-B (*t*(11) = −0.71, *p* = .508, *d* = −0.21, 95% CI [−0.26, 0.13]) showed a significant difference between conditions. In contrast, agentive movement elicited greater recruitment in DN-A (*t*(11) = −3.22, *p* = .018, *d* = −0.93, 95% CI [−0.51, −0.10]), DN-B (*t*(11) = −4.37, *p* = .005, *d* = −1.26, 95% CI [−0.67, −0.22]), and LANG (*t*(11) = −6.58, *p* < .001, *d* = −1.90, 95% CI [−1.04, −0.52]). Neither FPN-A (*t*(11) = −1.97, *p* = .096, *d* = −0.57, 95% CI [−0.48, 0.03]) nor FPN-B (*t*(11) = −2.41, *p* = .052, *d* = −0.70, 95% CI [−0.41, −0.02]) reached significance after FDR correction.

To summarize, networks associated with the canonical dorsal attention system (PM-PPr, dATN-A, and dATN-B) all showed above-baseline responses to both agentive and physical movement. Bilateral PM-PPr and CG-OP, and left dATN-A responded more strongly to physical movement. In contrast, dATN-B, which includes distributed regions in posterior parietal and occipitotemporal cortex, did not significantly differentiate between the two conditions. Frontoparietal control networks, which are often conflated with the dorsal attention system, showed a distinct response profile. Whereas networks associated with the canonical dorsal attention system generally preferred physical movement, FPN-A and FPN-B showed preferential responses to agentive movement in the left hemisphere, with effects in the same direction in the right hemisphere, although they did not survive FDR correction. These results highlight the importance of examining individual networks rather than treating broader system-level labels as monolithic.

Within the canonical default mode system, DN-B and LANG showed a preference for agentive movement across both hemispheres, while DN-A showed a significant agentive preference in the right hemisphere only. The preferential response of DN-B to agentive movement is consistent with its established role in social cognition and theory of mind, suggesting that this network may contribute specifically to interpreting actions performed by animate agents (DiNicola et al., 2020). The preference for the agentive condition only in the right DN-A is less straightforward to interpret and may reflect its spatial proximity to DN-B in association cortex. As for the LANG network, this network has been shown to overlap closely with regions involved in language processing (Braga et al., 2020; Du et al., 2024) and can be dissociated within individuals from the juxtaposed DN-A and DN-B networks in terms of its functional properties. The preferential responses to the agentive condition observed in the LANG network may reflect participants’ spontaneous linguistic or conceptual interpretation of the agentive interactions, such as labeling or describing events (e.g., “chasing” or “helping”), as the events depicted in the agentive condition naturally invited such interpretations. That said, because LANG, DN-B, and DN-A networks are highly interdigitated across much of the association cortex, it remains possible that some vertices attributed to one network may partially reflect signal from its juxtaposed neighbors due to spatial blurring.

## Discussion

In this study, we examined how neural responses to physical and agentive movement are spatially organized with respect to each other and in the context of functional network architecture. Several key findings emerged. First, a task requiring the interpretation and prediction of movement trajectories recruited a largely overlapping set of regions spanning frontoparietal and posterior temporal cortices, regardless of whether these predictions were based on the analysis of physical movement properties (physical condition) or inferences about agents and their intentions (agentive condition). At the same time, preferential responses for physical versus agentive movement dynamics were distributed throughout frontal, parietal, and posterior temporal cortices, often appearing in anatomically nearby but functionally dissociable regions. By integrating task-evoked responses with resting-state functional connectivity, we further demonstrated that brain regions preferentially responsive to physical movement were coupled at rest, as were regions preferring agentive movement.

### Network-level distinctions in processing physical versus agentive movement

Our work extends an emerging framework that distinguishes brain regions that support processing mechanical aspects of movement from those related to animate agents (Fischer et al., 2016; Isik et al., 2017; Jack et al., 2013; Martin & Weisberg, 2003; Mitchell et al., 2002; Pitcher & Ungerleider, 2021). However, previous accounts have largely focused on focal regional dissociations, leaving open how such functional distinctions are embedded within the brain’s intrinsic network organization.

Building on this literature and our previous work, we propose that a network largely overlapping with parts of the canonical dorsal attention system supports the analysis and interpretation of the physical and kinematic aspects of a dynamic scene, independent of whether the moving entity is animate or inanimate. In contrast, a distinct network overlapping with parts of the canonical default mode system preferentially supports computations that are specific to agents, such as inferring intentions or goals. Although the former network is also recruited when interpreting and predicting the movements of animate agents, agent-specific features appear to be preferentially represented within the latter network (for a similar thesis, see Karakose-Akbiyik et al., 2023; Liu et al., 2025).

This organization also bears on the proposed ‘third visual pathway,’ which locates dynamic social perception in a lateral occipitotemporal route centred on pSTS (Pitcher & Ungerleider, 2021). Here, regions around pSTS/TPJ that preferred agentive movement were embedded within a broader functional network that extended into prefrontal and posterior medial cortex, suggesting that this pathway operates within a more distributed system for interpreting agentive dynamics.

The proposal of two distinct networks, one for general kinematic reasoning and another for processes specific to animate agents, raises intriguing questions. In daily life, we integrate our understanding of physical dynamics with knowledge about animate agents to make meaningful inferences. To predict that someone will walk towards the kitchen because they seem thirsty, we must combine our understanding of the person’s needs with a physical prediction of their movement. If the first network supports predictions about physical dynamics, and the second encodes information specific to animate agents, does the latter provide input to the former? Conversely, we sometimes use our understanding of physical dynamics to infer how to interact with someone or to gauge their mental states. Observing someone reaching for an item on a high shelf, we assess their physical constraints or stability to determine whether they need help; from the physical costs of their actions, we infer their intentions and desires. This raises a further question: do these networks distinguish between physical information as it applies to animate agents versus inanimate objects?

Although these two modes of reasoning are often studied separately, they must work in concert, and an integrated account of how they converge is still needed. Future work could manipulate the relative emphasis on physical versus agentive dynamics and examine how representations within these networks change when reasoning must be integrated rather than separable (for similar ideas, see Liu et al., 2025). Techniques with higher temporal resolution, such as MEG, could further reveal when in the course of processing these two sources of information are combined.

Our findings also highlight that these functional distinctions do not map onto the broad network labels commonly used in the literature. Recent work using individual-subject analyses has demonstrated that large-scale networks such as the default mode and dorsal attention systems comprise multiple subsystems with distinct connectivity and task response profiles (Braga & Buckner, 2017; DiNicola et al., 2020; Kong et al., 2019, 2021). Consistent with this view, we observed involvement of specific network subdivisions in processing physical or agentive movement. For instance, within the so-called dorsal attention system, PM-PPr showed a bilateral preference for the physical condition, dATN-A showed this preference in the left hemisphere only, and dATN-B, which spans regions in posterior parietal and occipitotemporal cortices, did not differentiate the conditions in either hemisphere. CG-OP, which lies adjacent to the dorsal attention system and has been linked to action planning and physical inference (Badke D’Andrea et al., 2025; Dosenbach et al., 2025; Navarro-Cebrián & Fischer, 2022), also showed a bilateral preference for physical movement.

Fine-grained heterogeneity of this kind was particularly evident in lateral occipitotemporal cortex (LOTC), a region whose role in action recognition and physical reasoning has often been underappreciated. Lesion studies implicate anterior LOTC in action recognition, where damage produces impairments in recognizing observed actions (Tarhan et al., 2015; Urgesi et al., 2014). LOTC also hosts cross-exemplar and cross-modal representations of actions and exhibits a gradient from exemplar-specific to exemplar-general representations along its posterior–anterior axis (Lingnau & Downing, 2015; Wurm & Caramazza, 2019). A complementary organization is observed along its dorsal–ventral axis, with dorsal aspects preferentially engaged by bodies, biological motion, and person-directed actions, and ventral aspects preferentially engaged by tool motion and object-directed actions (Wurm & Caramazza, 2022). Finally, clusters in anterior LOTC are consistently reported in studies of physical and mechanical reasoning, although they sometimes exhibit functional properties that differ from those of frontoparietal regions implicated in the same tasks (Fischer et al., 2016; Pramod et al., 2022).

Our network analyses revealed that an anterior LOTC cluster was embedded within the same network as frontoparietal regions preferentially engaged by physical movement. Notably, this cluster did not always show the same preference as the frontoparietal components of this network, for instance, in some subjects, it did not show a preference for physical or agentive movement even though it was recruited for both (see Supplementary Figure 4). This suggests that while anterior LOTC is part of the same functional network that is involved in the analysis of physical and kinematic aspects of a dynamic scene, its specific contributions to this process may differ from those of frontoparietal cortices. Posterior LOTC, in contrast, fell outside this network and, in some participants, showed clusters preferring agentive movement. Together, these findings highlight that LOTC is not a homogeneous structure but comprises multiple subregions that participate in different networks and likely support complementary aspects of dynamic scene processing. This heterogeneity may help explain why LOTC has been difficult to place within existing frameworks of action and physical reasoning and underscores the value of network-based approaches.

### Animacy as an organizing principle for dynamic scene processing

Animacy is a critical dimension in the neural representation of object information and is increasingly recognized for its relevance in the context of processing dynamic information, particularly regarding action targets (e.g., person-vs. object-directed, Isik et al., 2017; Lee Masson & Isik, 2021; Wurm & Caramazza, 2019; Wurm & Schubotz, 2018). Building on this work, it has been suggested that the animacy-based organization found in the ventral temporal cortex for object recognition might also be present in the lateral temporal cortex for dynamic information processing (Wurm & Caramazza, 2022). Our findings extend this perspective by showing that anatomically nearby clusters in the frontal and parietal cortices also show preferences for aspects of dynamic scenes that are broadly associated with animate or inanimate entities.

More broadly, the correspondence between task-evoked preferences and intrinsic functional connectivity echoes findings from the ventral temporal cortex, where functional or structural connectivity recapitulates the organization of category-selective responses (Konkle & Caramazza, 2017; Osher et al., 2016; Saygin et al., 2012). In the context of ventral temporal and lateral occipitotemporal cortex and object recognition, functional connectivity predicts both the spatial layout and the representational geometry of object categories. Our results suggest a similar principle for the analysis of dynamic information: regions that preferentially respond to physical or agentive dynamics of movement are also embedded within distinct networks, indicating that the organization of dynamic-scene processing is similarly scaffolded by intrinsic connectivity patterns that reflect core dimensions such as animacy or agency.

It is important to note that we addressed only one aspect of variation by animacy, kinematic differences defined by Newtonian mechanics versus intentional, agent-driven movement. However, dynamic information processing involves a complex suite of component processes. For instance, processing the movement of an animate entity requires the analysis of self-propelled, agentive movement dynamics and biological motion, as well as higher-level processes related to intentions, goals, and social interactions. Conversely, analyzing the physical aspects of movement can involve processes such as inferring physical properties from movement (e.g., weight), predicting trajectories based on past movement, or encoding properties of objects for object-directed actions. These processes are often studied under the broad umbrella of intuitive physical reasoning, but the cognitive and neural mechanisms underlying them might be dissociable (Liu et al., 2026). In the current study, we observed multiple functional networks associated with both physical and agentive movement; however, this overlap does not imply that all these regions contribute in the same way to these component processes. Instead, they may support distinct functional components that were not isolated by our current design. Characterizing the specific computational roles of these networks remains an important objective for future research.

In summary, our study provides new insights into the distributed neural responses to physical and agentive movement dynamics. We find distributed preferences for physical and agentive movement across frontoparietal and posterior temporal cortices. These preferences were reflected in resting-state network architecture: regions with similar response profiles for physical or agentive movement exhibited correlated neural activity even at rest. Our findings extend existing frameworks of dynamic information processing and highlight the importance of considering connectivity between different brain regions in studying their functional organization.

## Materials and Methods

### Participants

Twelve right-handed participants (six female, *M*_age_ = 25.33, *SD*_age_ = 4.90) with normal or corrected-to-normal vision attended two scanning sessions, spaced at least a week apart. Before participating in the study, each participant provided informed consent. The experimental protocol was approved by Harvard University’s Committee on the Use of Human Subjects.

### DOTS task

Participants watched 32 unique 10-second movies featuring two dots. In the physical condition, dot movements followed Newtonian mechanics, changing paths due to collisions and forces, such as bouncing off walls. Conversely, in the agentive condition, the dots depicted coordinated self-propelled actions of animate agents (Fischer et al., 2016). In each movie, both dots moved around for 8 seconds, at which point one of them disappeared and reappeared after 2 seconds. The participants were prompted to mentally track its trajectory until it reappeared. Upon reappearance, they pressed a button to judge whether the dot returned to a plausible location. The physical and agentive conditions were matched for difficulty in terms of the trajectory judgment (see Fischer et al., 2016). Overall, participants were able to identify whether the dot reappeared at the right location (*M_physical_* = .82, *SD_physical_* = .07; *M_agentive_* = .78, *SD_agentive_* = .10) with no significant difference in accuracy (*t*(11) = 1.36, *p* = .20, *d* = .39, 95% CI [-.02, .09]) between the physical and agentive conditions.

Each run comprised 19 blocks of 26 seconds each: 8 blocks of physical movies, 8 of agentive movies, and 3 rest blocks showing a black screen. The blocks followed a palindromic sequence, in which pairs of physical and agentive blocks alternated, interspersed with rest blocks (e.g., A-A-P-P-A-A). In each non-rest block, participants viewed two consecutive movies of the same type (agentive or physical). After each 10-second movie, the final frame remained visible for 1.5 seconds for participants to judge and respond to the final dot position. A 1.5-second blank screen interval preceded the next movie. This sequence was repeated once more within the block, totaling 26 seconds per block. Rest blocks were placed at the beginning, middle, and end of the scanning run (blocks 1, 10, and 19), also organized in a palindromic pattern to maintain a structured flow in the experiment.

Note that in the original depiction of this task and while using it as a localizer, previous work collected two runs per participant (Fischer et al., 2016; Liu et al., 2024a). In the current study, each participant completed at least five runs of the DOTS task. This more intensive sampling was employed to boost signal-to-noise and to allow precise characterization of the organization of responses within individuals.

### Fixation runs

Participants were presented with a black crosshair against a light gray background. They were instructed to keep their eyes open, maintain fixation, and stay alert. No specific task was required. These runs served as the basis for the analysis of intrinsic functional connectivity. Each fixation run lasted 422 seconds (422 volumes with a 1-second TR). Each participant provided at least five fixation runs to allow estimation of individualized network parcellations and within-individual analyses of functional connectivity (Kong et al., 2019; Du et al., 2024).

### Data acquisition

Data were acquired at the Harvard Center for Brain Science using a 3T Siemens Prisma-fit MRI scanner. T1-weighted structural images were obtained using a 3D MPRAGE sequence (voxel size = 1 mm isotropic, TR = 2530 ms, TE = 1.69, 3.55, 5.41, and 7.27 ms, FOV: 256 x 256 mm, flip angle 7, 176 slices). For the acquisition of the BOLD signal, we used a custom multiband gradient-echo echo-planar pulse sequence developed by the Center for Magnetic Resonance Research (CMRR) at the University of Minnesota (voxel size = 2.4 mm isotropic, TR = 1,000 ms, TE = 33 ms, flip-angle = 64, (AP) encoding, matrix 92 x 92 x 65, FOV: 221 x 221mm, 65 slices).

### Data preprocessing

Functional and anatomical data were preprocessed using the publicly available *iProc* pipeline, a pipeline developed for single-subject datasets acquired over multiple sessions (Braga et al., 2019; Du et al., 2024). For all functional runs, we removed the first 12 frames to allow for T1 equilibration. We then corrected for head motion and field inhomogeneities, registered each run to a within-individual mean BOLD template, and aligned it to the participant’s T1-weighted anatomical image. For resting-state fixation runs, nuisance regressors comprising whole-brain, ventricular, and deep cerebral white matter signals, six head-motion parameters, and the temporal derivative of each were regressed out, and the residual data were band-pass filtered (0.01-0.10 Hz).

All preprocessed functional runs were projected to the fsaverage6 cortical surface using FreeSurfer and smoothed with a 2-mm full-width-at-half-maximum Gaussian kernel. These steps were implemented via a series of transformations that were concatenated and applied in a single interpolation step. Prior to the analysis of the fMRI data, the BOLD runs were inspected for quality. Exclusion criteria were adapted from Du et al. (2024) and included a maximum absolute head motion threshold of 2.5 mm and a slice-based SNR threshold of 130. Runs with an SNR between 100 and 130 were retained if they met the motion criterion and visual inspection of the scans displayed satisfactory quality. As a result of these criteria, a total of four resting-state runs and six runs of the DOTS task were discarded. All analyses were conducted on the fsaverage6 cortical surface and results were visualized using wb_view from the Connectome Workbench.

### Analysis of the DOTS task

The analysis of the DOTS task was conducted using the general linear model (GLM) via FSL’s first-level FEAT. The task followed a blocked design, with alternating blocks of physical and agentive movement trials modeled using a boxcar function convolved with a canonical double-gamma hemodynamic response function. Temporal derivatives were also included to account for HRF variability across brain regions. Low-frequency drifts were removed using high-pass temporal filtering with a 100-second cutoff (0.01 Hz). For each participant, z-value maps from all runs were combined to produce a single map for each contrast for each participant.

### Analysis of intrinsic functional connectivity

We examined intrinsic functional connectivity using both model-free seed-based and parcellation-based analyses. Initially, seed-based analyses were conducted to determine the extent to which task-related neural response organization is reflected in the brain’s intrinsic architecture. For these analyses, we calculated the pairwise Pearson correlation coefficients between the fMRI time courses of each surface vertex for each resting-state fixation run, resulting in an 81,924 x 81,924 matrix (40,962 vertices per hemisphere). We then transformed these matrices using Fisher’s r-to-z transformation and averaged the matrices across all runs to yield a single best estimate of the within-individual correlation matrix. To ensure reliable and consistent seed identification across participants, seeds were chosen for the two conditions in different hemispheres (left for physical; right for agentive), reflecting the lateralization of peak task preferences. Note that the resulting intrinsic connectivity maps for both seeds were bilateral and largely symmetrical (see Figures 4-5, Supplementary Figures 3 and 5), and all subsequent analyses evaluated networks in both hemispheres.

To characterize functional network responses to different task conditions, we also employed a seed-free approach using the multi-session hierarchical Bayesian model (MS-HBM) (Kong et al., 2019) as implemented in Du et al. (2024). This model begins with a group prior derived from Human Connectome Project S900 and generates individualized network maps by accounting for both session-to-session and person-to-person variabilities in functional connectivity profiles (Du et al., 2024). This framework enables the precise estimation of cortical parcellations for individual subjects, even with a limited amount of data collected over multiple sessions. The model identifies 15 networks: Somatomotor-A (SMOT-A), Somatomotor-B (SMOT-B), Premotor-Posterior Parietal Rostral (PM-PPr), Cingulo-Opercular (CG-OP), Salience/Parietal Memory Network (SAL/PMN), Dorsal Attention-A (dATN-A), Dorsal Attention-B (dATN-B), Frontoparietal Network-A (FPN-A), Frontoparietal Network-B (FPN-B), Default Network-A (DN-A), Default Network-B (DN-B), Language (LANG), Visual-Central (VIS-C), Visual-Peripheral (VIS-P), and Auditory (AUD).

## Data availability

Individual-participant contrast maps, network parcellations, and source data for Figures 6 and 8 will be deposited at the Open Science Framework (https://osf.io/zdmsh) and made publicly available upon publication.

## Competing interests

The authors declare no competing interests.

## Notes

https://osf.io/zdmsh

### Competing Interest Statement

The authors have declared no competing interest.

### Summary of Updates

This version adds a short Discussion section considering how the two proposed networks, one for physical/kinematic reasoning, one for agent-specific inference, may interact, and notes future directions (manipulating physical vs. agentive emphasis; MEG for timing). The remaining edits are minor clarifications of wording. No results, analyses, figures, or references were changed or removed.

## References

Adolphs, R. (2009). The social brain: neural basis of social knowledge. Annual Review of Psychology, 60(1), 693–716.

Albertini, D., Lanzilotto, M., Maranesi, M., & Bonini, L. (2021). Largely shared neural codes for biological and nonbiological observed movements but not for executed actions in monkey premotor areas. Journal of Neurophysiology, 126(3), 906–912.

Amodio, D. M., & Frith, C. D. (2006). Meeting of minds: the medial frontal cortex and social cognition. Nature Reviews. Neuroscience, 7(4), 268–277.

Andrews-Hanna, J. R., Reidler, J. S., Huang, C., & Buckner, R. L. (2010a). Evidence for the default network’s role in spontaneous cognition. Journal of Neurophysiology, 104(1), 322–335.

Andrews-Hanna, J. R., Reidler, J. S., Sepulcre, J., Poulin, R., & Buckner, R. L. (2010b). Functional-anatomic fractionation of the brain’s default network. Neuron, 65(4), 550– 562.

Badke D’Andrea, C., Laumann, T. O., Newbold, D. J., Lynch, C. J., Hadji, M., Nelson, S. M., Nielsen, A. N., Chauvin, R. J., Krimmel, S. R., Snyder, A. Z., Marek, S., Greene, D. J., Raichle, M. E., Dosenbach, N. U. F., & Gordon, E. M. (2025). Action-mode subnetworks for decision-making, action control, and feedback. Proceedings of the National Academy of Sciences of the United States of America, 122(27), e2502021122.

Beauchamp, M. S., & Martin, A. (2007). Grounding object concepts in perception and action: Evidence from FMRI studies of tools. Cortex; a Journal Devoted to the Study of the Nervous System and Behavior, 43(3), 461–468.

Braga, R. M., & Buckner, R. L. (2017). Parallel interdigitated distributed networks within the individual estimated by intrinsic functional connectivity. Neuron, 95(2), 457–471.e5.

Braga, R. M., DiNicola, L. M., Becker, H. C., & Buckner, R. L. (2020). Situating the left-lateralized language network in the broader organization of multiple specialized large-scale distributed networks. Journal of Neurophysiology, 124(5), 1415–1448.

Braga, R. M., Van Dijk, K. R. A., Polimeni, J. R., Eldaief, M. C., & Buckner, R. L. (2019). Parallel distributed networks resolved at high resolution reveal close juxtaposition of distinct regions. Journal of Neurophysiology, 121(4), 1513–1534.

Buckner, R. L., Andrews-Hanna, J. R., & Schacter, D. L. (2008). The brain’s default network: anatomy, function, and relevance to disease. Annals of the New York Academy of Sciences, 1124(1), 1–38.

Buckner, R. L., & DiNicola, L. M. (2019). The brain’s default network: updated anatomy, physiology and evolving insights. Nature Reviews. Neuroscience, 20(10), 593–608.

Buschman, T. J., & Kastner, S. (2015). From behavior to neural dynamics: An integrated theory of attention. Neuron, 88(1), 127–144.

Caspers, S., Zilles, K., Laird, A. R., & Eickhoff, S. B. (2010). ALE meta-analysis of action observation and imitation in the human brain. NeuroImage, 50(3), 1148–1167.

Corbetta, M., Patel, G., & Shulman, G. L. (2008). The reorienting system of the human brain: from environment to theory of mind. Neuron, 58(3), 306–324.

Corbetta, M., & Shulman, G. L. (2002). Control of goal-directed and stimulus-driven attention in the brain. Nature Reviews. Neuroscience, 3(3), 201–215.

Deen, B., & Freiwald, W. A. (2025). Parallel systems for social and spatial cognition reaching the cortical apex. Proceedings of the National Academy of Sciences of the United States of America, 122(44), e2520067122.

Deen, B., Koldewyn, K., Kanwisher, N., & Saxe, R. (2015). Functional organization of social perception and cognition in the superior temporal sulcus. Cerebral Cortex (New York, N.Y.: 1991), 25(11), 4596–4609.

Deen, B., Schwiedrzik, C. M., Sliwa, J., & Freiwald, W. A. (2023). Specialized networks for social cognition in the primate brain. Annual Review of Neuroscience, 46, 381–401.

DiNicola, L. M., Braga, R. M., & Buckner, R. L. (2020). Parallel distributed networks dissociate episodic and social functions within the individual. Journal of Neurophysiology, 123(3), 1144–1179.

Dixon, M. L., De La Vega, A., Mills, C., Andrews-Hanna, J. R., Spreng, R. N., Cole, M. W., & Christoff, K. (2018). Heterogeneity within the frontoparietal control network and its relationship to the default and dorsal attention networks. Proceedings of the National Academy of Sciences of the United States of America, 115(7), E1598–E1607.

Dosenbach, N. U. F., Fair, D. A., Miezin, F. M., Cohen, A. L., Wenger, K. K., Dosenbach, R. A. T., Fox, M. D., Snyder, A. Z., Vincent, J. L., Raichle, M. E., Schlaggar, B. L., & Petersen, S. E. (2007). Distinct brain networks for adaptive and stable task control in humans. Proceedings of the National Academy of Sciences of the United States of America, 104(26), 11073–11078.

Dosenbach, N. U. F., Raichle, M. E., & Gordon, E. M. (2025). The brain’s action-mode network. Nature Reviews. Neuroscience, 26(3), 158–168.

Du, J., DiNicola, L. M., Angeli, P. A., Saadon-Grosman, N., Sun, W., Kaiser, S., Ladopoulou, J., Xue, A., Yeo, B. T. T., Eldaief, M. C., & Buckner, R. L. (2024). Organization of the human cerebral cortex estimated within individuals: networks, global topography, and function. Journal of Neurophysiology, 131(6), 1014–1082.

Fedorenko, E., Ivanova, A. A., & Regev, T. I. (2024). The language network as a natural kind within the broader landscape of the human brain. Nature Reviews. Neuroscience, 25(5), 289–312.

Fischer, J., Mikhael, J. G., Tenenbaum, J. B., & Kanwisher, N. (2016). Functional neuroanatomy of intuitive physical inference. In Proceedings of the National Academy of Sciences (Vol. 113, Issue 34, pp. E5072–E5081). 10.1073/pnas.1610344113

Fox, M. D., Corbetta, M., Snyder, A. Z., Vincent, J. L., & Raichle, M. E. (2006). Spontaneous neuronal activity distinguishes human dorsal and ventral attention systems. Proceedings of the National Academy of Sciences of the United States of America, 103(26), 10046– 10051.

Fox, M. D., Snyder, A. Z., Vincent, J. L., Corbetta, M., Van Essen, D. C., & Raichle, M. E. (2005). The human brain is intrinsically organized into dynamic, anticorrelated functional networks. Proceedings of the National Academy of Sciences of the United States of America, 102(27), 9673–9678.

Gazzola, V., & Keysers, C. (2009). The observation and execution of actions share motor and somatosensory voxels in all tested subjects: single-subject analyses of unsmoothed fMRI data. Cerebral Cortex (New York, N.Y.: 1991), 19(6), 1239–1255.

Gilaie-Dotan, S., Kanai, R., Bahrami, B., Rees, G., & Saygin, A. P. (2013). Neuroanatomical correlates of biological motion detection. Neuropsychologia, 51(3), 457–463.

Gordon, E. M., Laumann, T. O., Gilmore, A. W., Newbold, D. J., Greene, D. J., Berg, J. J., Ortega, M., Hoyt-Drazen, C., Gratton, C., Sun, H., Hampton, J. M., Coalson, R. S., Nguyen, A. L., McDermott, K. B., Shimony, J. S., Snyder, A. Z., Schlaggar, B. L., Petersen, S. E., Nelson, S. M., & Dosenbach, N. U. F. (2017). Precision functional mapping of individual human brains. Neuron, 95(4), 791–807.e7.

Grèzes, J., Fonlupt, P., Bertenthal, B., Delon-Martin, C., Segebarth, C., & Decety, J. (2001). Does perception of biological motion rely on specific brain regions? NeuroImage, 13(5), 775–785.

Grossman, E. D., & Blake, R. (2002). Brain areas active during visual perception of biological motion. Neuron, 35(6), 1167–1175.

Hardwick, R. M., Caspers, S., Eickhoff, S. B., & Swinnen, S. P. (2018). Neural correlates of action: Comparing meta-analyses of imagery, observation, and execution. Neuroscience and Biobehavioral Reviews, 94, 31–44.

Heilman, K. M., & Valenstein, E. (1972). Frontal lobe neglect in man. Neurology, 22(6), 660– 664.

Isik, L., Koldewyn, K., Beeler, D., & Kanwisher, N. (2017). Perceiving social interactions in the posterior superior temporal sulcus. Proceedings of the National Academy of Sciences of the United States of America, 114(43), E9145–E9152.

Jack, A. I., Dawson, A. J., Begany, K. L., Leckie, R. L., Barry, K. P., Ciccia, A. H., & Snyder, A. Z. (2013). fMRI reveals reciprocal inhibition between social and physical cognitive domains. NeuroImage, 66, 385–401.

Karakose-Akbiyik, S., Caramazza, A., & Wurm, M. F. (2023). A shared neural code for the physics of actions and object events. Nature Communications, 14(1), 1–13.

Karakose-Akbiyik, S., Sussman, O., Wurm, M. F., & Caramazza, A. (2024). The Role of Agentive and Physical Forces in the Neural Representation of Motion Events. The Journal of Neuroscience: The Official Journal of the Society for Neuroscience, 44(2). 10.1523/JNEUROSCI.1363-23.2023

Kastner, S., & Ungerleider, L. G. (2000). Mechanisms of visual attention in the human cortex. Annual Review of Neuroscience, 23, 315–341.

Kennedy, D. P., & Adolphs, R. (2012). The social brain in psychiatric and neurological disorders. Trends in Cognitive Sciences, 16(11), 559–572.

Kim, M. J., Maione, S., & Liu, S. (2026). Shared neural representations of barriers to agents’ actions and objects’ motions. In PsyArXiv. 10.31234/osf.io/v2dgf_v1

Koldewyn, K., Whitney, D., & Rivera, S. M. (2011). Neural correlates of coherent and biological motion perception in autism. Developmental Science, 14(5), 1075–1088.

Kong, R., Li, J., Orban, C., Sabuncu, M. R., Liu, H., Schaefer, A., Sun, N., Zuo, X.-N., Holmes, A. J., Eickhoff, S. B., & Yeo, B. T. T. (2019). Spatial topography of individual-specific cortical networks predicts human cognition, personality, and emotion. Cerebral Cortex (New York, N.Y.: 1991), 29(6), 2533–2551.

Kong, R., Yang, Q., Gordon, E., Xue, A., Yan, X., Orban, C., Zuo, X.-N., Spreng, N., Ge, T., Holmes, A., Eickhoff, S., & Yeo, B. T. T. (2021). Individual-specific areal-level parcellations improve functional connectivity prediction of behavior. Cerebral Cortex (New York, N.Y.: 1991), 31(10), 4477–4500.

Konkle, T., & Caramazza, A. (2017). The large-scale organization of object-responsive cortex is reflected in resting-state network architecture. *Cerebral Cortex (New York*, N.Y.: 1991). 10.1093/cercor/bhw287

Laufs, H., Krakow, K., Sterzer, P., Eger, E., Beyerle, A., Salek-Haddadi, A., & Kleinschmidt, A. (2003). Electroencephalographic signatures of attentional and cognitive default modes in spontaneous brain activity fluctuations at rest. Proceedings of the National Academy of Sciences of the United States of America, 100(19), 11053–11058.

Lee Masson, H., & Isik, L. (2021). Functional selectivity for social interaction perception in the human superior temporal sulcus during natural viewing. NeuroImage, 245(118741), 118741.

Lieberman, M. D. (2007). Social cognitive neuroscience: a review of core processes. Annual Review of Psychology, 58(1), 259–289.

Lingnau, A., & Downing, P. E. (2015). The lateral occipitotemporal cortex in action. Trends in Cognitive Sciences, 19(5), 268–277.

Liu, S., Karakose-Akbiyik, S., Caramazza, A., Dourado Sotero, F., Buxbaum, L. J., & Wong, A. L. (2026). Dissociation between physical reasoning and tool use in individuals with left hemisphere brain damage. Cortex; a Journal Devoted to the Study of the Nervous System and Behavior, 201, 148–170.

Liu, S., Karakose-Akbiyik, S., Outa, J., & Kim, M. J. (2025). How physical information is used to make sense of the psychological world. Nature Reviews Psychology. 10.1038/s44159-025-00514-1

Liu, S., Lydic, K., Mei, L., & Saxe, R. (2024a). Violations of physical and psychological expectations in the human adult brain. Imaging Neuroscience, 2, 1–25.

Liu, S., Wurm, M. F., & Caramazza, A. (2024b). Dissociating goal from outcome during action observation. *Cerebral Cortex (New York*, N.Y.: 1991), *34*(12). 10.1093/cercor/bhae487

Mars, R. B., Neubert, F.-X., Noonan, M. P., Sallet, J., Toni, I., & Rushworth, M. F. S. (2012). On the relationship between the “default mode network” and the “social brain.” Frontiers in Human Neuroscience, 6, 189.

Martin, A., & Weisberg, J. (2003). Neural foundations for understanding social and mechanical concepts. Cognitive Neuropsychology, 20(3–6), 575–587.

Meyer, M. L. (2019). Social by default: Characterizing the social functions of the resting brain. Current Directions in Psychological Science, 28(4), 380–386.

Mitchell, J. P., Heatherton, T. F., & Macrae, C. N. (2002). Distinct neural systems subserve person and object knowledge. Proceedings of the National Academy of Sciences of the United States of America, 99(23), 15238–15243.

Navarro-Cebrián, A., & Fischer, J. (2022). Precise functional connections between the dorsal anterior cingulate cortex and areas recruited for physical inference. The European Journal of Neuroscience, 56(1), 3660–3673.

Osher, D. E., Saxe, R. R., Koldewyn, K., Gabrieli, J. D. E., Kanwisher, N., & Saygin, Z. M. (2016). Structural connectivity fingerprints predict cortical selectivity for multiple visual categories across cortex. Cerebral Cortex (New York, N.Y.: 1991), 26(4), 1668–1683.

Pitcher, D., & Ungerleider, L. G. (2021). Evidence for a Third Visual Pathway Specialized for Social Perception. Trends in Cognitive Sciences, 25(2), 100–110.

Pramod, R. T., Cohen, M. A., Tenenbaum, J. B., & Kanwisher, N. (2022). Invariant representation of physical stability in the human brain. eLife, 11. 10.7554/eLife.71736

Ptak, R. (2012). The frontoparietal attention network of the human brain: action, saliency, and a priority map of the environment. *The Neuroscientist: A Review Journal Bringing Neurobiology*, Neurology and Psychiatry, 18(5), 502–515.

Ptak, R., Schnider, A., & Fellrath, J. (2017). The dorsal frontoparietal network: A core system for emulated action. Trends in Cognitive Sciences, 21(8), 589–599.

Raichle, M. E. (2015). The brain’s default mode network. Annual Review of Neuroscience, 38(1), 433–447.

Raichle, M. E., MacLeod, A. M., Snyder, A. Z., Powers, W. J., Gusnard, D. A., & Shulman, G. L. (2001). A default mode of brain function. Proceedings of the National Academy of Sciences of the United States of America, 98(2), 676–682.

Rushworth, M., Walton, M., Kennerley, S., & Bannerman, D. (2004). Action sets and decisions in the medial frontal cortex. Trends in Cognitive Sciences, 8(9), 410–417.

Saygin, A. P., Wilson, S. M., Hagler, D. J., Jr, Bates, E., & Sereno, M. I. (2004). Point-light biological motion perception activates human premotor cortex. The Journal of Neuroscience: The Official Journal of the Society for Neuroscience, 24(27), 6181–6188.

Saygin, Z. M., Osher, D. E., Koldewyn, K., Reynolds, G., Gabrieli, J. D. E., & Saxe, R. R. (2012). Anatomical connectivity patterns predict face selectivity in the fusiform gyrus. Nature Neuroscience, 15(2), 321–327.

Scholl, B. J., & Gao, T. (2013). Perceiving Animacy and Intentionality. In Social Perception (pp. 197–230). The MIT Press.

Schurz, M., Maliske, L., & Kanske, P. (2020). Cross-network interactions in social cognition: A review of findings on task related brain activation and connectivity. Cortex; a Journal Devoted to the Study of the Nervous System and Behavior, 130, 142–157.

Schurz, M., Radua, J., Aichhorn, M., Richlan, F., & Perner, J. (2014). Fractionating theory of mind: a meta-analysis of functional brain imaging studies. Neuroscience and Biobehavioral Reviews, 42, 9–34.

Schwettmann, S., Tenenbaum, J. B., & Kanwisher, N. (2019). Invariant representations of mass in the human brain. eLife, 8. 10.7554/eLife.46619

Simone, L., Pierotti, E., Satta, E., Becchio, C., & Turella, L. (2025). Resting-state functional interactions between the action observation network and the mentalizing system. The European Journal of Neuroscience, 61(6), e70082.

Smith, V., Mitchell, D. J., & Duncan, J. (2018). Role of the default mode network in cognitive transitions. Cerebral Cortex (New York, N.Y.: 1991), 28(10), 3685–3696.

Tarhan, L., Watson, C. E., & Buxbaum, L. J. (2015). Shared and distinct neuroanatomic regions critical for tool-related action production and recognition: Evidence from 131 left-hemisphere stroke patients. Journal of Cognitive Neuroscience, 27(12), 2491–2511.

Urgesi, C., Candidi, M., & Avenanti, A. (2014). Neuroanatomical substrates of action perception and understanding: an anatomic likelihood estimation meta-analysis of lesion-symptom mapping studies in brain injured patients. Frontiers in Human Neuroscience, 8, 344.

Vallar, G., & Perani, D. (1987). The Anatomy of Spatial Neglect in Humans. In M. Jeannerod (Ed.), Advances in Psychology (Vol. 45, pp. 235–258). North-Holland.

Van Overwalle, F., & Baetens, K. (2009). Understanding others’ actions and goals by mirror and mentalizing systems: a meta-analysis. NeuroImage, 48(3), 564–584.

Wang, Y., Metoki, A., Alm, K. H., & Olson, I. R. (2018). White matter pathways and social cognition. Neuroscience and Biobehavioral Reviews, 90, 350–370.

Wurm, M. F., & Caramazza, A. (2019). Distinct roles of temporal and frontoparietal cortex in representing actions across vision and language. Nature Communications, 10(1), 289.

Wurm, M. F., & Caramazza, A. (2022). Two “what” pathways for action and object recognition. Trends in Cognitive Sciences, 26(2), 103–116.

Wurm, M. F., & Erigüç, D. Y. (2025). Decoding the physics of observed actions in the human brain. eLife, 13(RP98521). 10.7554/elife.98521.3

Wurm, M. F., & Schubotz, R. I. (2018). The role of the temporoparietal junction (TPJ) in action observation: Agent detection rather than visuospatial transformation. NeuroImage, 165, 48–55.

Yan, Y., Zhan, J., Garrod, O. G. B., Ince, R. A. A., Jack, R. E., & Schyns, P. G. (2025). The brain computes dynamic facial movements for emotion categorization using a third pathway. Proceedings of the National Academy of Sciences of the United States of America, 122(25), e2423560122.

Yeo, B. T. T., Krienen, F. M., Sepulcre, J., Sabuncu, M. R., Lashkari, D., Hollinshead, M., Roffman, J. L., Smoller, J. W., Zöllei, L., Polimeni, J. R., Fischl, B., Liu, H., & Buckner, R. L. (2011). The organization of the human cerebral cortex estimated by intrinsic functional connectivity. Journal of Neurophysiology, 106(3), 1125–1165.

Yeshurun, Y., Nguyen, M., & Hasson, U. (2021). The default mode network: where the idiosyncratic self meets the shared social world. Nature Reviews. Neuroscience, 22(3), 181–192.

Ziccarelli, S., Errante, A., & Fogassi, L. (2022). Decoding point-light displays and fully visible hand grasping actions within the action observation network. Human Brain Mapping, 43(14), 4293–4309.

