## Supplementary Materials for "Spatial organization of neural responses to physical and agentive movement dynamics is reflected in intrinsic functional connectivity"

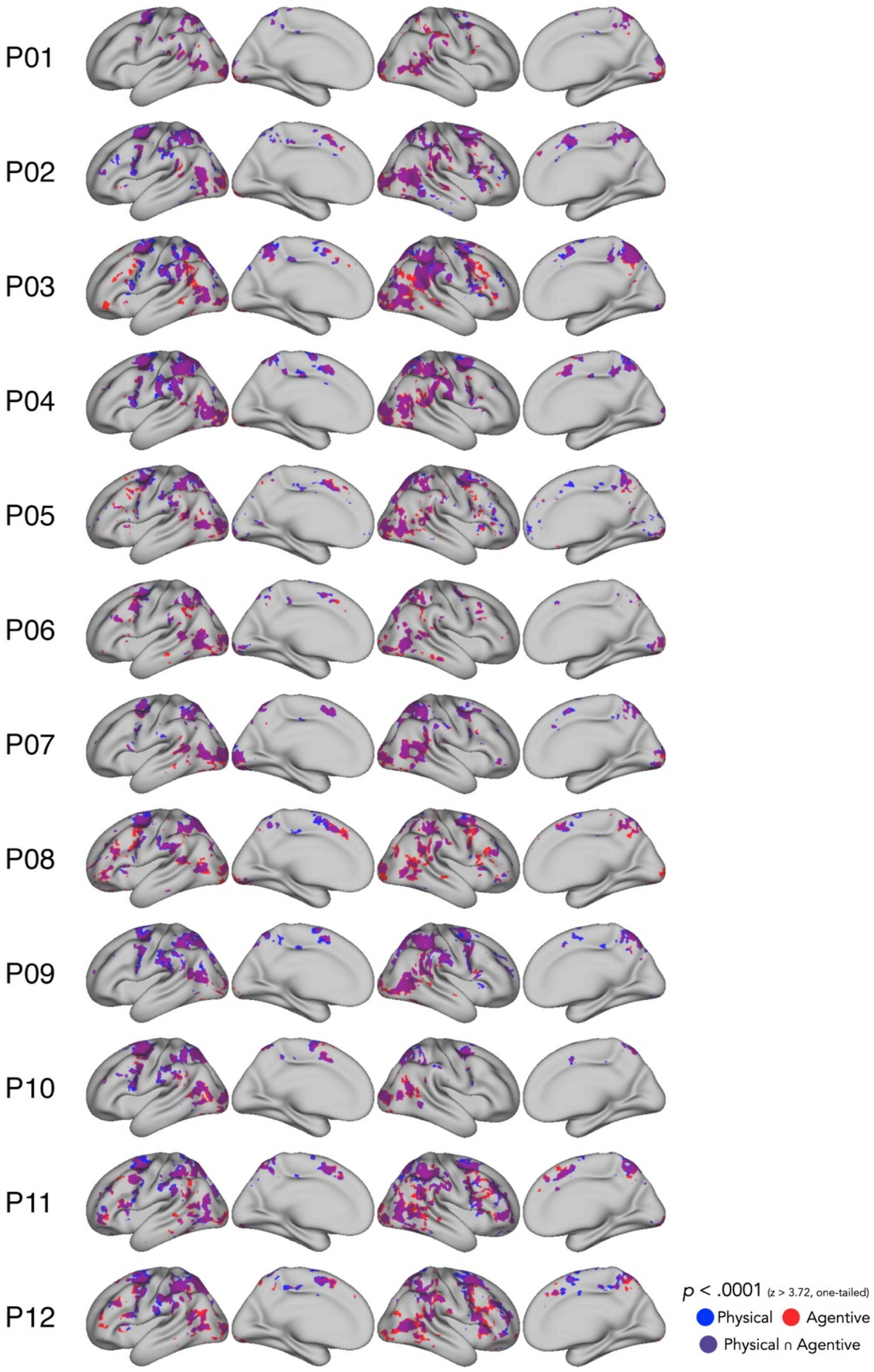
P01
P02
P03
P04
P05
P06
P07
P08
P09
P10
P11
P12
p < .0001 (z > 3.72, one-tailed)
Physical
Agentive
Physical ∩ Agentive

**Supplementary Figure 1. Neural responses to physical and agentive movement for all participants.** Brain regions with increased responses compared to baseline during the DOTS task for physical and agentive conditions. Blue areas represent regions with higher than baseline activity exclusively for physical movement, red areas for agentive movement, and purple areas show the overlap where both conditions elicited higher-than-baseline activity. Threshold: $p < .0001$, $z > 3.72$, one-tailed, uncorrected.

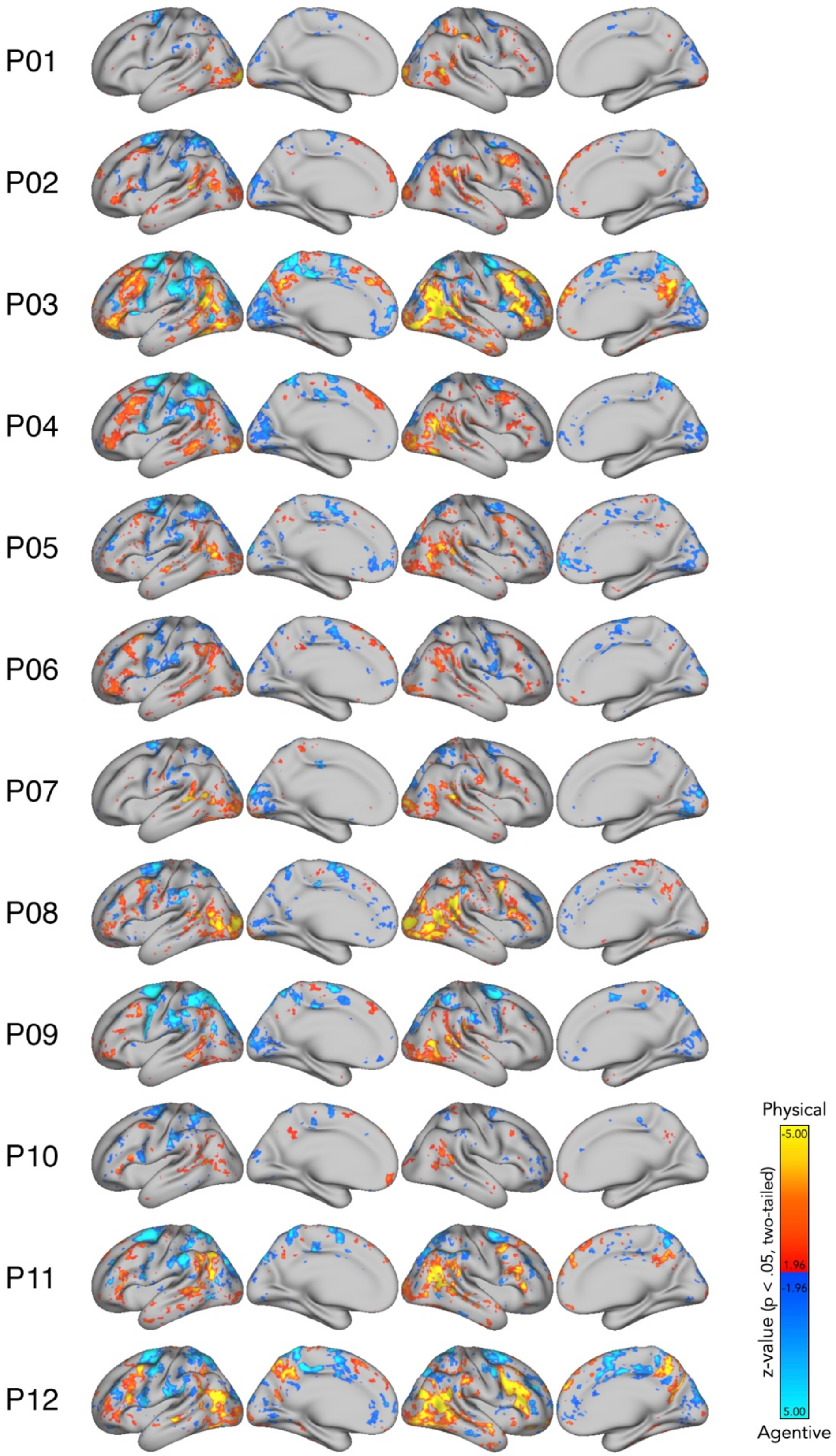
P01
P02
P03
P04
P05
P06
P07
P08
P09
P10
P11
P12
Physical
-5.00
1.96
-1.96
5.00
z-value (p < .05, two-tailed)
Agentive

**Supplementary Figure 2. Univariate contrast of agentive and physical movement in all participants.** Blue regions show greater responses to physical stimuli, while orange/yellow regions show greater responses to agentive stimuli. Maps are shown at a liberal threshold ($p < .05$, $|z| > 1.96$, two-tailed, uncorrected) to illustrate individual variability in response patterns.

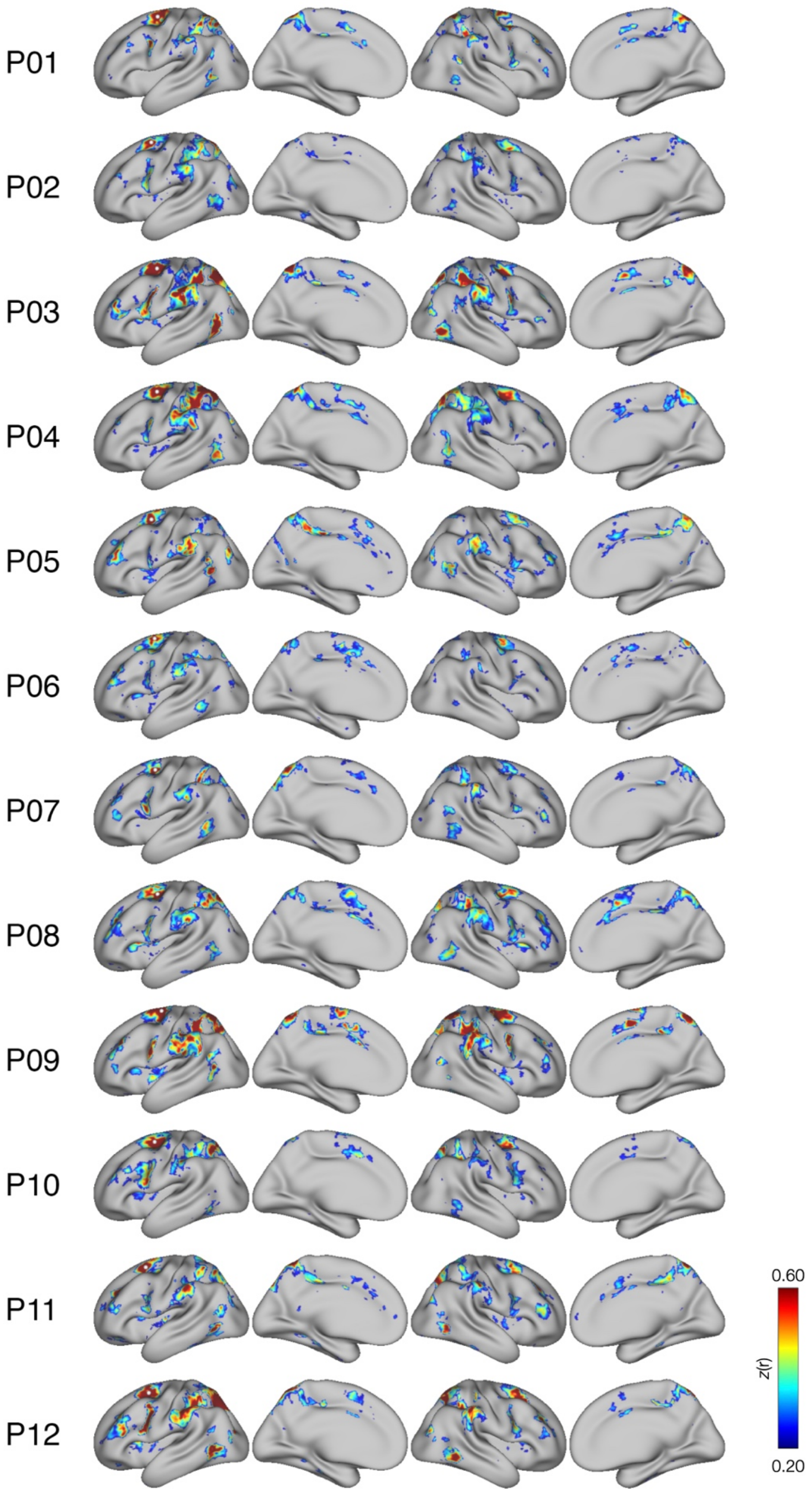
P01
P02
P03
P04
P05
P06
P07
P08
P09
P10
P11
P12
0.60
z(r)
0.20

**Supplementary Figure 3. Seed-based functional connectivity maps for all participants for the physical seed.** Functional connectivity maps for all participants using the seed chosen for the physical condition. The white dot marks the location of the seed used for the connectivity analysis. The maps were thresholded with $z$(r) values between 0.20 and 0.60.

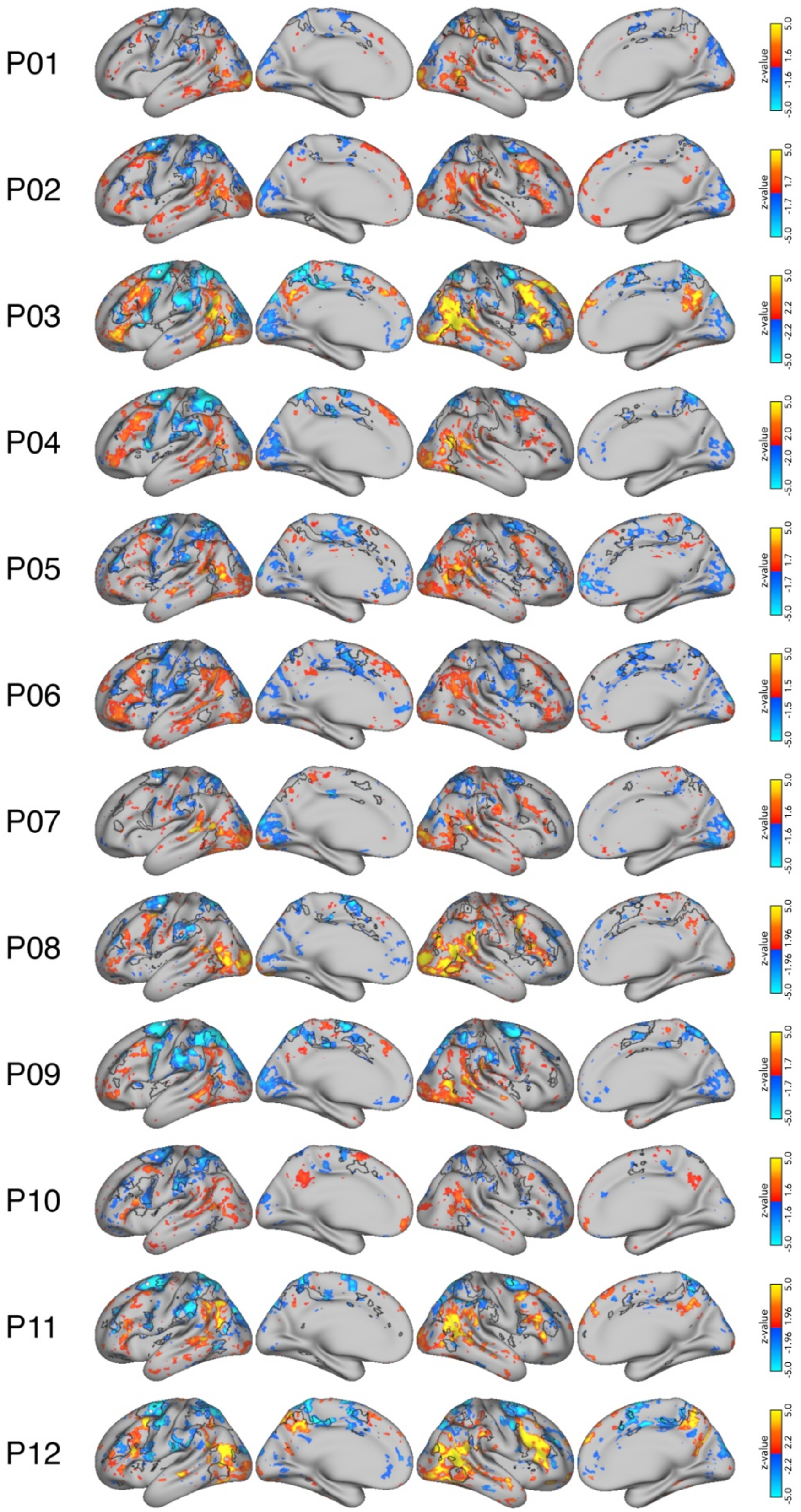
P01
P02
P03
P04
P05
P06
P07
P08
P09
P10
P11
P12
z-value
5.0
1.6
-1.6
-5.0
1.7
-1.7
2.2
-2.2
2.0
-2.0
1.5
-1.5
1.96
-1.96

**Supplementary Figure 4. Mapping preferential responses to physical and agentive movement to seed-based functional connectivity maps for the physical seed.** Overlay of the agentive – physical contrast maps in the DOTS task with the seed-based functional connectivity maps generated for the physical seed. The black outlines indicate the boundaries of the functional connectivity maps thresholded with *z*(r) values between 0.20 and 0.60. The DOTS task contrasts show regions with a preference for physical (blue) and agentive (orange/yellow) movement. For visualization, DOTS contrast maps were displayed using a liberal and varying threshold (|z| = 1.5-2.2) to better illustrate individual trends.

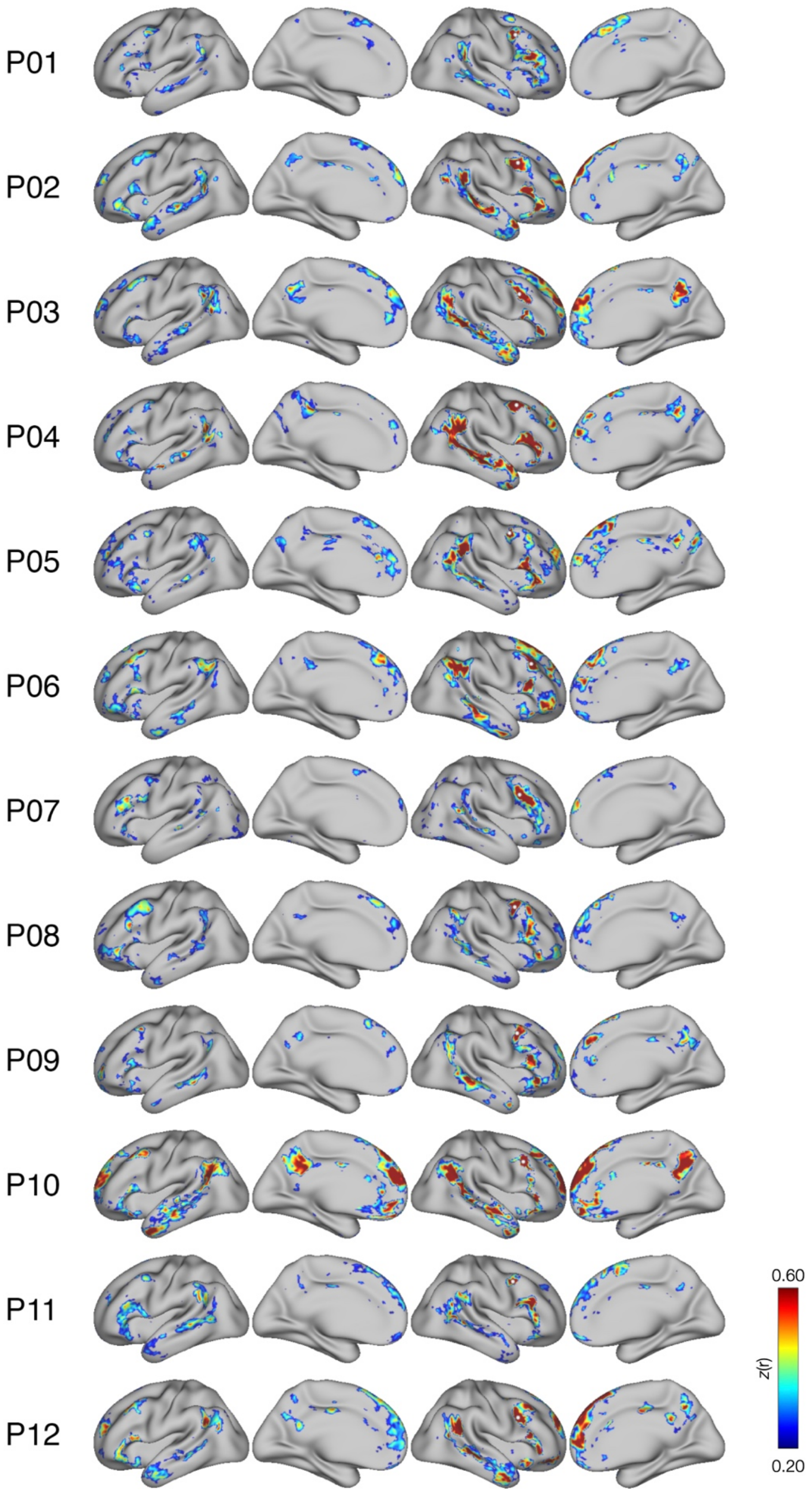
P01
P02
P03
P04
P05
P06
P07
P08
P09
P10
P11
P12
0.60
z(r)
0.20

**Supplementary Figure 5. Seed-based functional connectivity maps for all participants for the agentive seed.** Functional connectivity maps for all participants using the seed chosen for the agentive condition. The white dot marks the location of the seed used for the connectivity analysis. The maps were thresholded with *z*(r) values between 0.20 and 0.60.

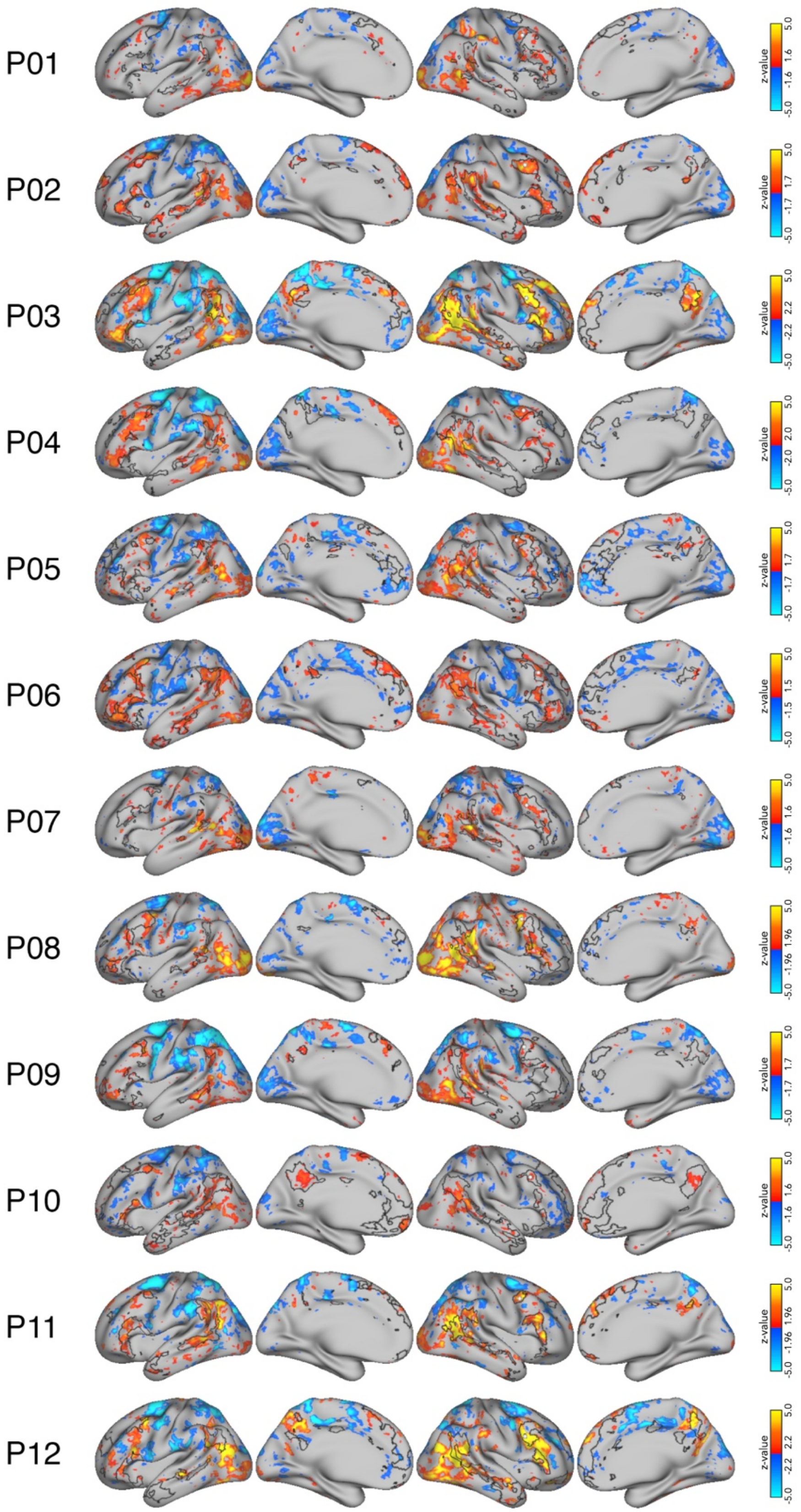
P01
z-value 5.0 1.6 -1.6 -5.0
P02
z-value 5.0 1.7 -1.7 -5.0
P03
z-value 5.0 2.2 -2.2 -5.0
P04
z-value 5.0 2.0 -2.0 -5.0
P05
z-value 5.0 1.7 -1.7 -5.0
P06
z-value 5.0 1.5 -1.5 -5.0
P07
z-value 5.0 1.6 -1.6 -5.0
P08
z-value 5.0 1.96 -1.96 -5.0
P09
z-value 5.0 1.7 -1.7 -5.0
P10
z-value 5.0 1.6 -1.6 -5.0
P11
z-value 5.0 1.96 -1.96 -5.0
P12
z-value 5.0 2.2 -2.2 -5.0

**Supplementary Figure 6. Mapping preferential responses to physical and agentive movement to seed-based functional connectivity maps for the agentive seed.** Overlay of the agentive – physical contrast maps in the DOTS task with the seed-based functional connectivity maps generated for the agentive seed. The black outlines indicate the boundaries of the functional connectivity maps thresholded with $z$(r) values between 0.20 and 0.60. The DOTS task contrasts show regions with a preference for physical (blue) and agentive (orange/yellow) movement. For visualization, DOTS contrast maps were displayed using a liberal and varying threshold ($|z| = 1.5$-$2.2$) to better illustrate individual trends.

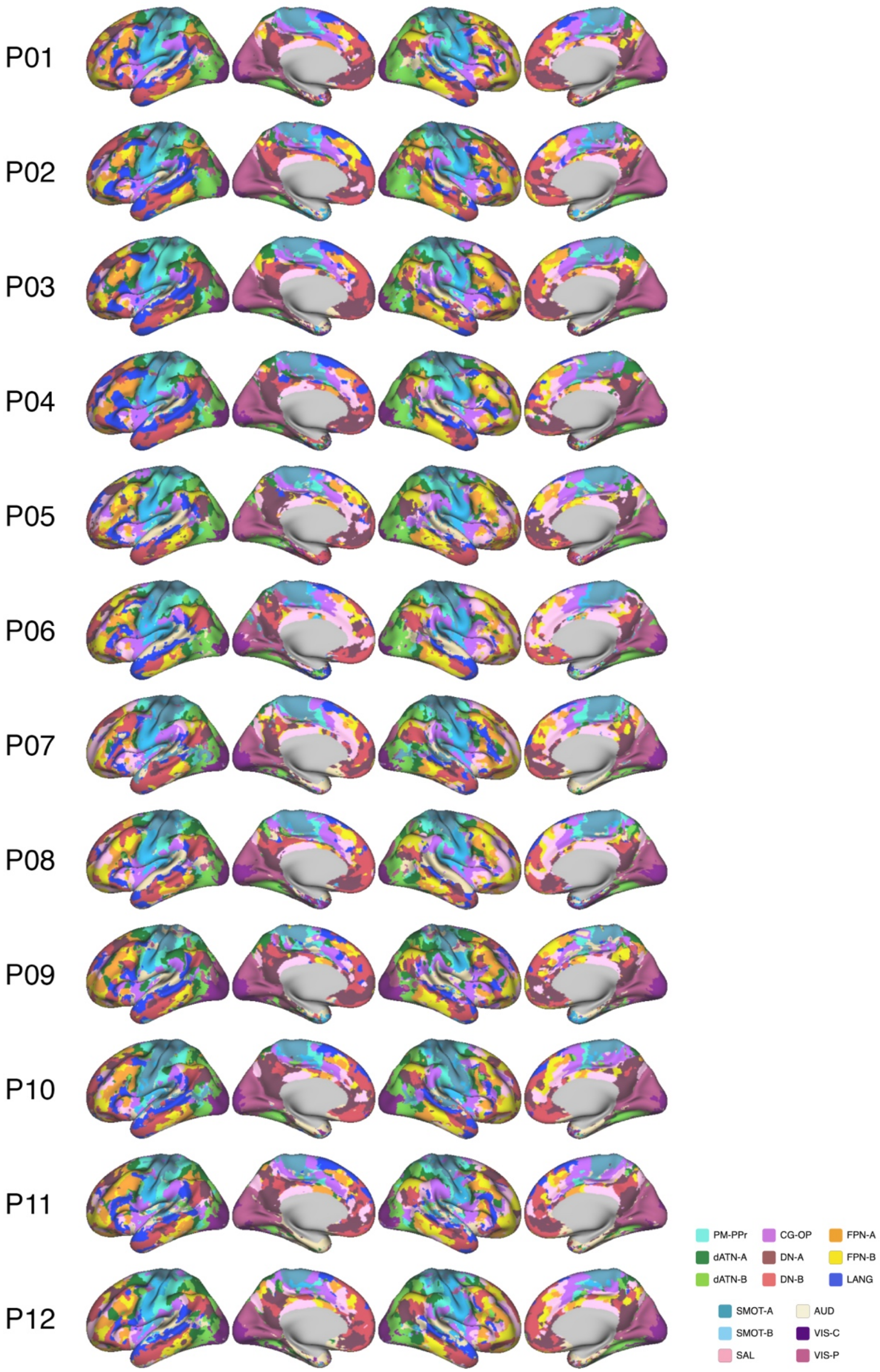
P01
P02
P03
P04
P05
P06
P07
P08
P09
P10
P11
P12
PM-PPr
CG-OP
FPN-A
dATN-A
DN-A
FPN-B
dATN-B
DN-B
LANG
SMOT-A
AUD
SMOT-B
VIS-C
SAL
VIS-P

**Supplementary Figure 7. 15-Network Parcellation for all participants.** 15-Network estimation for identifying the cortical networks for all participants. Cortical networks were estimated for each participant using a multi-session hierarchical Bayesian model, following the implementation described in Du et al. (2024). The model was initialized with a group prior generated from the Human Connectome Project S900.
